# Modulation of coordinated activity across cortical layers by plasticity of inhibitory synapses onto layer 5 pyramidal neurons

**DOI:** 10.1101/567065

**Authors:** Joana Lourenço, Angela Michela De Stasi, Charlotte Deleuze, Mathilde Bigot, Antonio Pazienti, Andrea Aguirre, Michele Giugliano, Srdjan Ostojic, Alberto Bacci

**Affiliations:** Sorbonne Universités UPMC Univ. Paris 06, UMR S 1127, Inserm U 1127, CNRS UMR 7225, and ICM-Institut du Cerveau et de la Moelle épinière, 75013 Paris, France; Laboratoire de Neurosciences Cognitives, INSERM U960, Ecole Normale Supérieure - PSL Research University, 75005 Paris, France; European Brain Research Institute, Fondazione Rita Levi-Montalcini, 00143 Rome, Italy; Department of Biomedical Sciences and Institute Born-Bunge, Molecular, Cellular, and Network Excitability, Universiteit Antwerpen, Antwerpen, Belgium

**Author notes:** Corresponding authors: Alberto Bacci; Joana Lourenço. Istituto Superiore di Sanità, 00161 Rome, Italy (Antonio Pazienti).

## Abstract

In the neocortex, synaptic inhibition shapes all forms of spontaneous and sensory-evoked activity. Importantly, inhibitory transmission is highly plastic, but the functional role of inhibitory synaptic plasticity is unknown. In the mouse barrel cortex, activation of layer 2/3 PNs elicited strong feed-forward perisomatic inhibition (FFI) onto layer 5 PNs. We found that FFI involving PV cells was strongly potentiated by postsynaptic PN burst firing. FFI plasticity modified PN excitation-to-inhibition (E/I) ratio, strongly modulated PN gain and altered information transfer across cortical layers. Moreover, our LTPi-inducing protocol modified the firing of layer 5 PNs and altered the temporal association of PN spikes to γ-oscillations both in vitro and in vivo. All these effects were captured by unbalancing the E/I ratio in a feed-forward inhibition circuit model. Altogether, our results indicate that activity-dependent modulation of perisomatic inhibitory strength effectively influences the participation of single principal cortical neurons to cognitive-relevant network activity.

**Impact Statement:** Long-term potentiation of feed-forward perisomatic inhibition effectively alters the computational properties of single layer 5 pyramidal neurons and their association to network activity.

## Introduction

In the neocortex, sensory integration is accomplished through the coordinated activity of neuronal networks across different cortical layers and columns (reviewed in Allene et al., 2015;Douglas et al., 2004;Feldmeyer, 2012). These hardwired anatomical connectivity patterns between several neuron types define specific pathways and enable a salient flow of information across and within different cortical layers.

Functional cortical networks result from both direct contact between neurons and by indirect feed-forward and feedback connections from intercalated neurons, whose recruitment strength and excitability contribute to the formation and dissolution of neuronal ensembles (Buzsaki, 2010). These interposed neurons are mostly (but not only) inhibitory. Importantly, fast synaptic inhibition plays a fundamental role in shaping and sculpting virtually all forms of cortical activity (Sachdev et al., 2012; Atallah et al., 2012;Lee et al., 2012;Wilson et al., 2012; Buzsaki and Wang, 2012;Cardin et al., 2009;Sohal et al., 2009;Veit et al., 2017).

In the neocortex, inhibition is provided by a rich diversity of GABAergic interneurons (Ascoli et al., 2008;Kepecs and Fishell, 2014;Mendez and Bacci, 2011). In particular, perisomatic targeting interneurons account for nearly half of all cortical interneurons and form inhibitory synapses on the cell body and proximal dendrites of their targets (Bodor et al., 2005;Freund and Katona, 2007). Inhibitory transmission alters the computations performed by PNs, strongly affecting their output activity (Carandini and Heeger, 2011;Silver, 2010). For example, in the visual cortex, it has been shown that perisomatic inhibition provided by PV cells controls the gain of visual responses, affecting PN sensitivity to sensory stimuli without changing their feature selectivity (Atallah et al., 2012;Lee et al., 2012;Wilson et al., 2012).

In addition, PV interneurons play a key role in the generation of fast cortical oscillations in the β-γ frequency range (20-100 Hz) (Cardin et al., 2009;Sohal et al., 2009; but see Veit et al., 2017), believed to underlie several cognitive functions, such as sensory perception and attention (Bartos et al., 2007;Buzsaki and Silva, 2012;Wang, 2010). PV cell-mediated perisomatic inhibition onto PNs thus acts as a synchronizing mechanism, locking the spike timing of a large population of PNs to a specific oscillation phase, and effectively promoting the generation of cell assemblies (Buzsaki, 2010).

We have previously shown that postsynaptic depolarization or bursts of action potentials in layer 5 PNs of the mouse barrel cortex induces a long-term potentiation of inhibition (LTPi), which is selective for PV cell-transmission and relies on a Ca_2+-_dependent retrograde signaling of nitric oxide (Lourenço et al., 2014). This non-associative potentiation of perisomatic GABAergic synapses strongly alters the excitation-to-inhibition (E/I) ratio on layer 5 PNs, reduces firing probability and sharpens the time window of synaptic integration (Lourenço et al., 2014).

Long-term plasticity of glutamatergic synapses has been the focus of intense investigation and it has been postulated to be the synaptic correlate of learning and memory (Malenka, 2003). Yet, despite the fact that inhibitory synaptic transmission is highly plastic (Castillo et al., 2011;Griffen and Maffei, 2014;Mendez and Bacci, 2011), the functional role of GABAergic plasticity is unknown (but see (Mongillo et al., 2018;Vogels et al., 2011)). Here, we set out to investigate how plasticity of PV cell-mediated perisomatic GABAergic synapses modulates several computations performed by single layer 5 PNs of the mouse barrel cortex (S1). Using in utero electroporation, we expressed light-sensitive opsins in layer 2/3 PNs of mouse S1. We demonstrate that activation of layer 2/3 induces robust feed-forward inhibition (FFI) on layer 5 PNs, mediated by PV basket cells. FFI could be strongly potentiated by cell-autonomous postsynaptic paradigms. LTPi of FFI modified input/ouput relationship of layer 5 PNs and prevented their increased excitability, induced by layer 2/3 activation. Moreover, LTPi-inducing bursts affected the temporal association of PN spiking with γ-oscillations both in vitro and in vivo. These results were captured by a computational model, indicating that plasticity of PV cell-dependent perisomatic inhibition can strongly modulate single PNs both at the single-cell and network level.

## Results

### Burst firing of layer 5 pyramidal neurons selectively potentiates feed-forward GABAergic inputs

Activity of layer 2/3 PNs, in addition to inducing lateral spreading via horizontal intralaminar connections, plays a prominent role in activating deep cortical output layers, and represents a prominent excitatory pathway onto large layer 5 PNs. We have previously demonstrated that bursts of action potentials (APs) in layer 5 PNs induce LTPi selectively at synapses from PV cells (Lourenço et al., 2014). We therefore asked whether layer 2/3 activation triggers feed-forward inhibition (FFI) onto layer 5 PNs, and whether FFI is plastic.

Using *in utero* electroporation, we expressed the light-sensitive opsin channelrhodospin 2 (ChR2) in a large fraction of layer 2/3 PNs (Figure 1A). We then performed whole-cell patch-clamp recordings from large layer 5 PNs in acute slices of barrel cortex (S1, barrel field) from mice, which had been previously electroporated *in utero* (Figure 1 A and B). Brief (1 ms) stimulations with blue light (λ = 470 nm) of layer 2/3 ChR2+ PNs induced a composite postsynaptic potential (PSP) in layer 5 PNs recorded in current clamp mode at resting membrane potential. This composite PSP was composed of an early excitatory postsynaptic potential (EPSP), followed by an inhibitory postsynaptic potential (IPSP; Figure 1B and 1C, top panel). This IPSP was likely triggered by perisomatic-targeting PV cells, as activation of layer 2/3 PNs recruited layer 5 PV interneurons efficiently (Figure 1 – figure supplement 1).

**Figure 1.**
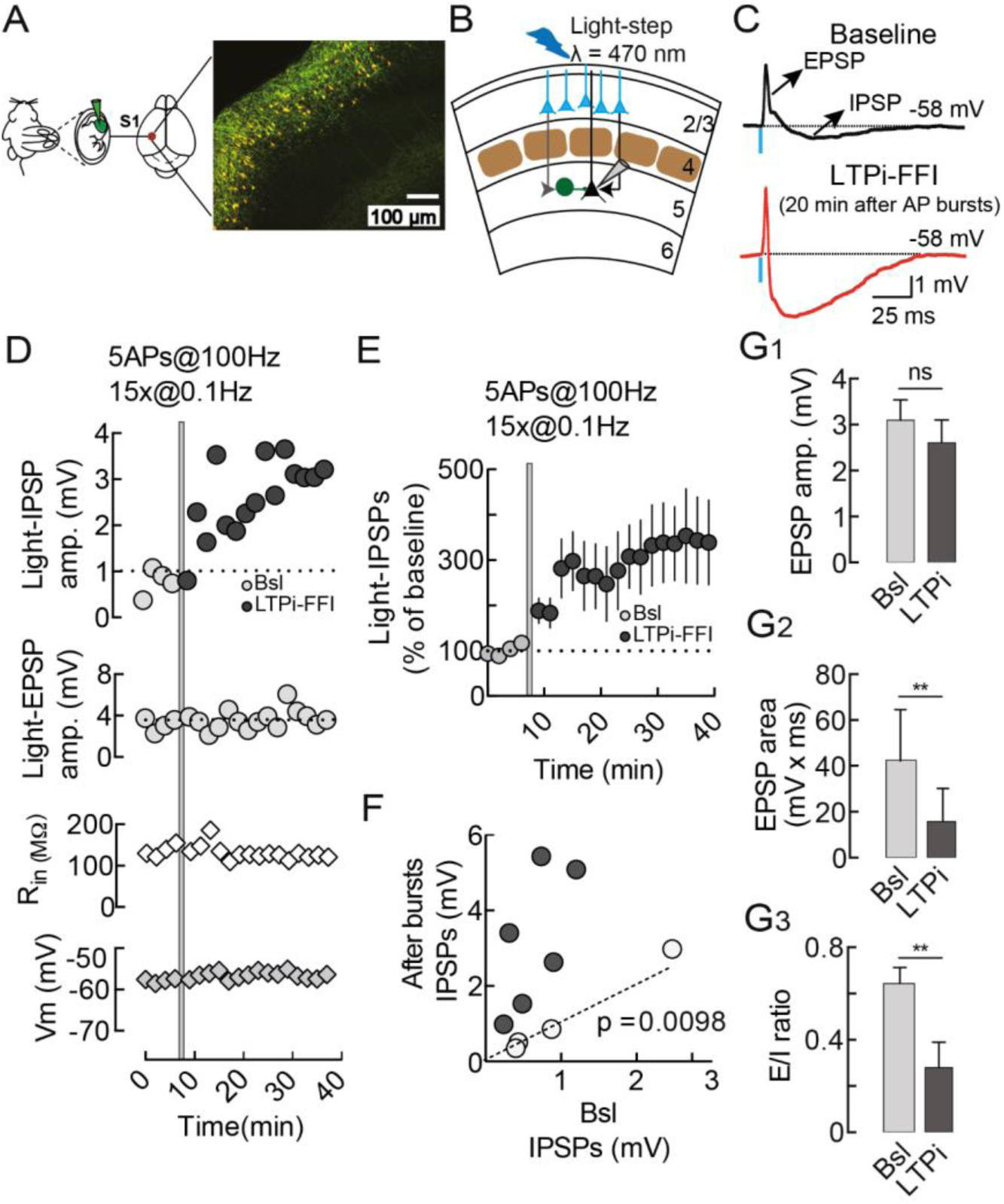
Burst firing of layer 5 pyramidal neurons selectively potentiates feed-forward GABAergic inputs. (**A**)*in utero* electroporation of ChR2 and the red fluorophore RFP in layer 2/3 PNs of the mouse somatosensory barrel cortex. (**B**) Scheme of the recording configuration of layer 5 PNs, stimulated by ChR2+ layer 2/3 PNs; black triangle represents the soma of layer 5 PNs and green full circle represents the soma of a PV cell. (**C**)Representative current-clamp trace of the EPSP-IPSPs composite recorded in layer 5 PN upon photostimulation of layer 2/3 PNs. Note the increase in the IPSP component (Red trace, LTPi) after postsynaptic bursts. Each trace is the average of 10 sweeps. (**D**) Time course of light-IPSPs (top graph) of the cell shown in (C), displaying a clear LTPi-FFI. The shaded area refers to the LTPi-FFI induction protocol - postsynaptic action potentials bursts. Light-EPSPs (graph below), input resistance (Rin, Middle) and resting membrane potential (Vm, Bottom) remained stable throughout the experiment. (**E**) Population time course of normalized light-IPSPs during baseline (gray circles) and after LTPi-FFI induction (black circles) averaged in 2 min bins. (**F**) Plot of individual eIPSC amplitudes before (x-axes) versus 20 min after postsynaptic bursts (y-axes). The majority of layer 5 PNs expressed a long-term change in light-eIPSP amplitude, which we designated as LTPi-FFI (grey circles). A small percentage of the cells do not express LTPi (open circles). Dotted line indicates unitary values (no change).Grey symbols and white symbols refer to pyramidal neurons that did and did not express LTPi, respectively. (**G**) Graphs showing average depolarizing peaks, areas, and EPSP/IPSP ratio of composite PSPs in baseline and after postsynaptic bursts. The peak of the depolarizing component was not changed after LTPi-FFI induction(G1). However, a significant reduction of EPSP area (G2) and EPSC/IPSC ratio (G3) was observed upon postsynaptic bursts. For (**E**) and (**G**) the error bars indicate SEM. In some cases, the error bars are too small tobe visible. n.s: not significant. *p<0.05, **p<0.01, with paired *t* test.

Importantly, in response to postsynaptic AP bursts of layer 5 PNs (5APs at 100Hz, repeated 15 times every 10 seconds), we observed an increase in amplitude of the light-evoked IPSP component (light-IPSP) that persisted for >30 minutes (Figure 1C-F), which we termed long-term potentiation of feed-forward inhibition (LTPi-FFI). LTPi-FFI was observed in ∼ 65% of the recorded PNs (Figure 1 – figure supplement 2A, includes all cells tested for LTPi-FFI in Figures 1, 2 and 4) and it induced an increase in light-IPSPs amplitude of at least 50% of baseline amplitude (0.73 ± 0.2 vs. 2.38 ± 0.58 mV, light-IPSP baseline vs. 20 min after AP bursts, respectively; n = 10, p = 0.0098, Wilcoxon matched-pairs signed rank test, Figure 1F and Figure 1 – figure supplement 2A). We previously reported that this LTPi is selective for PV-cell synapses and induced by PN AP bursts and due to a Ca_2+-_dependent retrograde signaling of nitric oxide (NO) (Lourenço et al., 2014). Likewise, LTPi-FFI was also sensitive to pharmacological inhibition of the canonical NO receptor guanylylcyclase (GC) with 1H-{1,2,4}oxadiazolo{4,3-a}quinoxalin-&-dione (ODQ, 10μM; 0.73 ± 0.16 vs. 0.89 ± 0.31 mV, light-IPSP baseline vs. 20 min after AP bursts, respectively, in the continuous presence of ODQ; n = 8, p = 0.74, Wilcoxon matched-pairs signed rank test; Figure 1 - figure supplement 2B and C). Importantly, LTPi-FFI did not affect the peak amplitude of the depolarizing EPSP component (Baseline, 3.096 ± 0.44 mV; LTPi, 2.606 ± 0.49 mV, n= 10, p= 0.1, paired *t* test; Figure 1D and G_1_). However, due to the potentiation of the hyperpolarizing (IPSP) component, the area of the light-evoked EPSP significantly decreased (Baseline: 42.46 ± 7.47 mV*ms; LTPi: 15.73 ± 4.8 mV*ms; n= 10, p= 0.0036, paired *t* test; Figure 1G_2_). Consequently, LTPi-FFI strongly reduced the excitation-to-inhibition (E/I) ratio, measured as the EPSP area divided by the total composite PSP area (0.64 ± 0.07 vs. 0.28 ± 0.11, baseline vs. LTPi; n= 10, p= 0.0023, paired *t* test; Figure 1G_3_). In the presence of the NO receptor inhibitor ODQ, postsynaptic AP bursts failed to induce changes in the E/I ratio (0.79 ± 0.28 vs. 0.636 ± 0.1 mV, baseline vs. 20 min after AP bursts, in the continuous presence of ODQ; n = 8, p = 0.12, Wilcoxon matched-pairs signed rank test; Figure 1 – figure supplement 2D).

**Figure 2.**
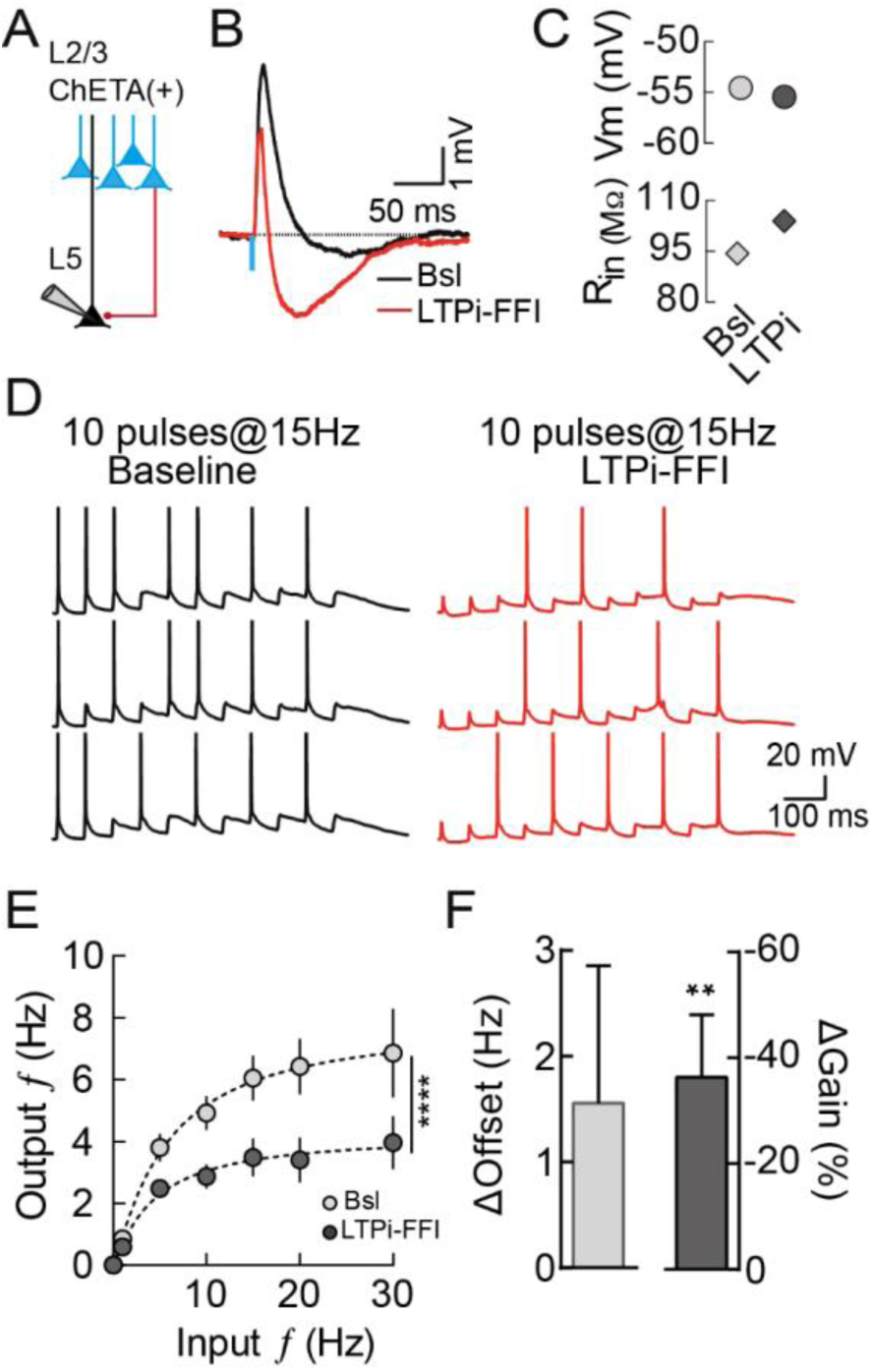
LTPi-FFI induces a multiplicative gain modulation of PN output spiking. (**A**) Schematic recording configuration of layer 5 PNs stimulated by ChETA+ layer 2/3 PNs (**B**) Representative current-clamp trace of the EPSP-IPSPs composite recorded in layer 5 PN upon photostimulation of ChETA+ layer 2/3 PNs. Note the increase in the IPSP component (Red trace, LTPi-FFI) after postsynaptic bursts. Each trace is the average of 10 sweeps. (**C**) Resting membrane potential (Vm, left panel) and input resistance (Rin, right panel) stable throughout the experiment. (**D**) Representative current-clamp traces of the firing output of layer 5 PN upon photostimulation of layer 2/3 PNs at 15Hz frequency during baseline and after LTPi-FFI (Red trace). Note the reduction in action potentials after LTPI-FFI (action potentials were clipped). (**E**) Graph illustrating averaged output firing rate of layer 5 PNs upon photostimulation of layer 2/3 PNs at different frequencies before (Bsl, light gray, n = 16) and 15 min after LTPi induction (LTPi-FFI, dark gray, n =16, ****p<0.0001, Friedman test). Dashed lines are fits to a Hill function (**F**) Offset (light gray) and gain (dark gray) changes in the input/ouput function after LTPi-FFI from fit in D (**p<0.01, Wilcoxon matched-pairs signed rank test, theoretical median = 0). For (**E**) and (**F**)the error bars indicate SEM.

In sum, PV cell-mediated FFI onto layer 5 PNs, triggered by layer 2/3 PN axons, can be strongly potentiated by postsynaptic firing activity alone. FFI plasticity resulted in prolonged changes of the E/I balance.

### LTPi-FFI induces a divisive gain modulation of PN output spiking

The input/output (I/O) relationship often reflects the way a neuron transforms input signals into output spike trains (Silver, 2010). Plastic changes of inhibitory synaptic strength might therefore modulate the ability of single PNs to encode input spike trains. To test this hypothesis, we measured the I/O relationship before and after inducing LTPi-FFI in layer 5 PNs by postsynaptic AP bursts (Figure 2A-C). To reliably evoke presynaptic spike trains, we expressed the fast light-sensitive opsin ChETA in layer 2/3 PNs via *in utero* electroporation. We then tested its effectiveness by recording spike activity in ChETA-expressing layer 2/3 PNs, in response to trains of brief blue light-pulses over a range of different frequencies (Figure 2 - figure supplement 1A-C). *In utero* electroporated layer 2/3 PNs linearly followed ChETA stimulations at frequencies up to 30 Hz (Figure 2 - figure supplement 1C). The effect of LTPi-FFI on synaptic integration were then investigated by stimulating layer 2/3 ChETA(+) neurons with 10 pulses of light at different frequencies, and measuring the mean output firing rate of layer 5 PNs (Figure 2D and E) before and after inducing LTPi-FFI. We observed an overall difference between baseline and LTPi-FFI with a major impact on higher frequencies between 15 and 30Hz (F_(11,165)_=13.76, p<0.0001; p<0.05 for 15 Hz, p<0.0001 for 20 Hz and p<0.05 for 30 Hz, Friedman test followed by Dunn’s multiple comparison test; Figure 2E). The input/output relationship was quantified by fitting the data to Hill-like equations, which allowed us extracting slope and offset of I-O curves (Murphy and Miller, 2003;Rothman et al., 2009). We observed a shift in the input frequency required to achieve half-maximum of the output frequency, although this was not significantly different (Δ_Offset_ = 1.56 ± 1.3 Hz, n = 14, p = 0.8, Wilcoxon matched-pairs signed rank test, theoretical median = 0; Figure 2F, light grey bar). Despite an absence of significant changes in the subtractive shift behavior of the input/output curve, we observed a significant change in the slope (Δ_Gain_ = −36.29 ± 11.79 %, n = 14, p = 0.01, Wilcoxon matched-pairs signed rank test, theoretical median = 0; Figure 2F, dark grey bar). Importantly, in the minority of layer 5 PNs that failed to express LTPi-FFI (Figure 1 – figure supplement 2A) we did not observe any change in gain modulation (Figure 2 – figure supplement 1D). Thus, these results indicate that LTPi-FFI has as an almost purely divisive effect on the PN input/output relationship (Figure 2E and F).

### LTPi-FFI affects the flow information across cortical layers

Sustained firing of layer 2/3 PNs induces lateral suppression of superficial cortical layers, and simultaneously activates layer 5 PNs within the same column. This latter phenomenon is known as feed-forward facilitation (FFF), and it is due to differences in E/I ratios in different cortical layers (Adesnik and Scanziani, 2010). Given the above-mentioned effects of LTPi-FFI on layer 5 PN E/I ratio and on the input/output function at the single cell level, we hypothesized that layer 2/3-induced facilitation of deep cortical layer PNs could be affected by potentiation of local perisomatic inhibitory transmission. FFF can be induced in cortical slices by depolarizing ChR2+ layer 2/3 PNs with a 1-s-long ramp of blue light, while simultaneously depolarizing layer 5 PNs (Adesnik and Scanziani, 2010). Layer 5 PNs were depolarized with 1-s-long current injections to trigger action potential firing (average rate: 5.4 ± 0.29 Hz, range 2–8 Hz, n = 23, Figure 3A and C, Bsl, black trace and grey bar), which was significantly increased by simultaneous photo-stimulation of layer 2/3 (average rate: 8.26 ± 0.35 Hz, range 5–12 Hz, n = 23, Figure 3A and C, blue trace and bar, Bsl_light; F(3,66)=35.83, p<0.0001; p<0.0001 for Bsl versus Bsl_light, Friedman test followed by Dunn’s multiple comparison test). Remarkably, when the same experiment was repeated after inducing LTPi-FFI, the facilitation of layer 5 PN excitability, triggered by layer 2/3 activation, was largely decreased (average rate: 6.22 ± 0.41 Hz, range 2–10 Hz, n = 23, Figure 3B and C red trace and red bar, LTPi_light; F(3,66)=35.83, p<0.0001; p = 0.7218 for LTPi vs. LTPi_light, Friedman test followed by Dunn’s multiple comparison test). It is known that FFF results from sustained firing of layer 2/3 PNs. Accordingly, LTPi-FFI did not change the baseline spike rate of layer 5 PNs (average rate 5.873 ± 0.35 Hz, range 3–8 Hz, n = 23, Figure 3B and C, black trace and dotted bar, LTPi), ruling out the possibility that postsynaptic burst firing alone altered layer 5 PN excitability.

**Figure 3.**
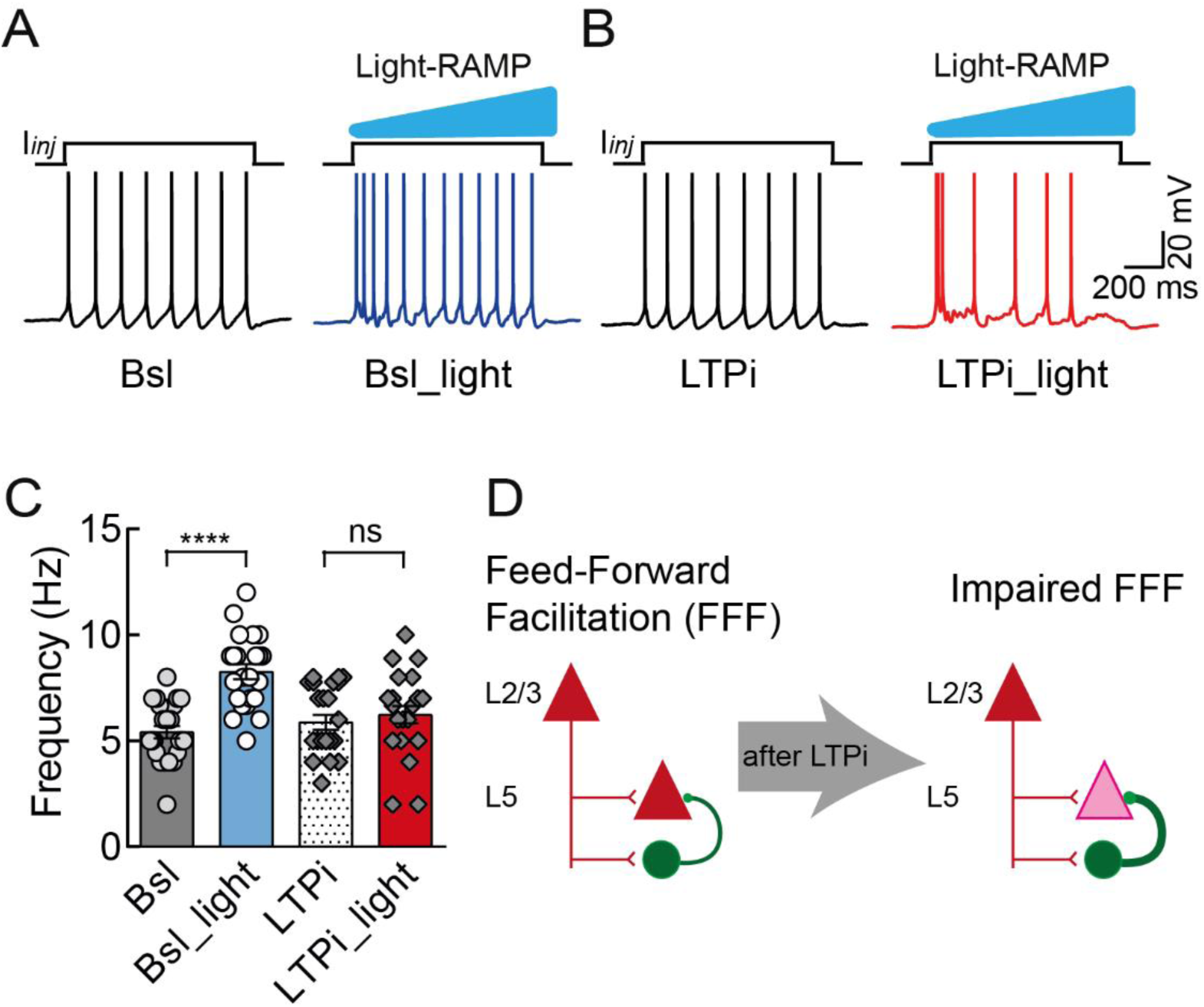
LTPi-FFI affects the flow information across cortical layers. (**A**) Response of a layer 5 PN to a depolarizing current injection (+70 pA), in baseline conditions without (left trace, Bsl) and with light stimulation (right trace, Bsl_light). Note the increase action potentials frequency (blue trace) upon photostimulation. The phenomenon of increase in spiking output of layer 5 upon stimulation of layer 2/3 PNs is known as feed-forward facilitation (FFF) (Adesnik and Scanziani, 2010). (**B**) Same cell of (A), upon LTPi-inducing AP bursts, in the absence (LTPi) and presence (LTPi_light) of light stimulation. Note that FFF of layer 5 PNs (red trace, light stimulation) is reduced upon LTPi-FFI. In (A) and (B) action potentials were clipped for display purposes. (**C**) Average population data of FFF during baseline condition in the absence and presence of light (circles) and upon LTPI-FFI (diamonds), ****p<0.0001, Friedman test. Error bars indicate SEM. (**D**) Scheme summarizing the effect of LTPi in preventing FFF across cortical layers. In the absence of LTPi, layer 2/3 activity efficiently spreads to layer 5 (left). Potentiation of feed-forward inhibition prevents this prominent information transfer from layer 2/3 to layer 5 PNs.

**Figure 4.**
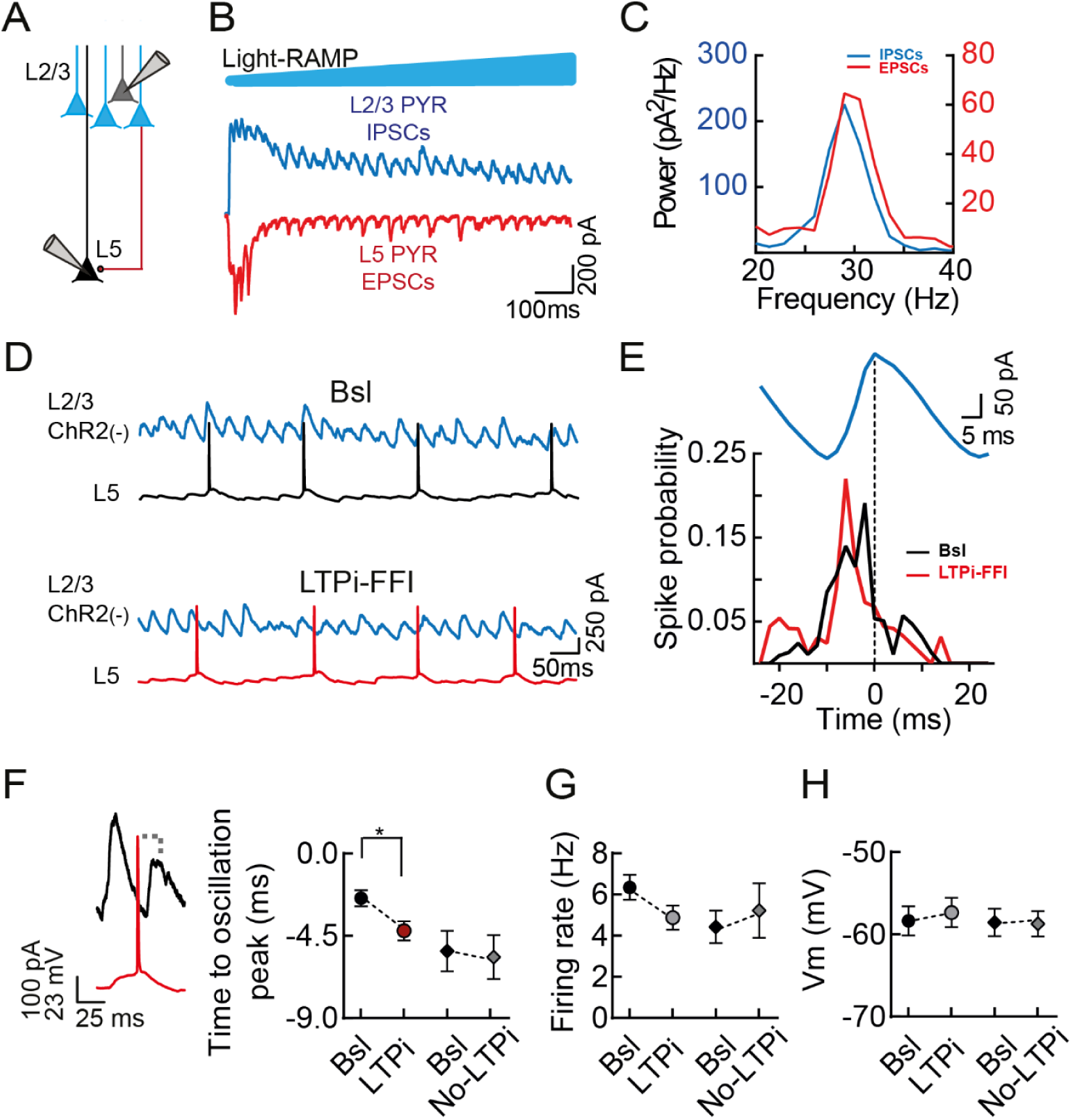
LTPi-FFI induces a shift in temporal association of layer 5 PN firing during photo-induced rhythmic activity. (**A**) Scheme of the recording configuration: ChR2-negative layer 2/3 and layer 5 PNs were recorded, while stimulating ChR2+ layer 2/3 PNs (blue) with by blue light. (**B**) Representative traces during a light ramp (blue, 1-s duration) protocol. The ramp photostimulation generated rhythmic activity of layer 2/3 IPSCs (recorded in voltage-clamp at +10mV, blue trace) and of layer 5 EPSCs (recorded in voltage-clamp at −70mV, red trace). (**C**) Power spectrum of the EPSCs and IPSCs of the cells shown in (**B**). (**D**) Schematic recording configuration and representative traces of the same cells as in (**B**). In order to access the temporal association of layer 5 PNs to rhythmic activity of layer 2/3 IPSCs (recorded in voltage-clamp at +10mV, blue trace) we recorded layer 5 in current-clamp configuration during baseline and after LTPi-FFI (black and red traces). (**E**) Spike probability during a rhythmic cycle (blue trace, recorded in layer 2/3 PNs) of the same layer 5 PN during baseline (black line) after LTPi-FFI (red line). (**F**) Left panel: representative traces illustrating the association of a spike of a layer 5 PN (red trace) with the peak of oscillating IPSCs from a layer 2/3 PN (black trace). Right panel: Average population data of the time to oscillation peak in cells that underwent LTPI-FFI (circles, n = 9) and cells, in which the postsynaptic bursts had no effect (diamonds, n =6), n = 11, *p<0.05, Wilcoxon matched-pairs signed rank test. (**G**) Average population data of the firing rate of LTPI-FFI (circles) and no LTPi cells (diamonds). (**H**) Average population data of the membrane resting potentials of LTPI-FFI (circles) and no LTPi cells (diamonds). In (D) and (F) action potentials were clipped. For (**F**), (**G**) and (**H**) the error bars indicate SEM.

Altogether, these results indicate that the cell-autonomous long-term strengthening of perisomatic inhibition can reduce or completely abolish the coordination of cortical activity across cortical layers (Figure 3D).

### LTPi-FFI induces a shift in temporal association of layer 5 PN firing during photo-induced rhythmic activity

One of the most prominent functional roles of perisomatic inhibition, particularly from PV basket cells, is that of synchronizing large populations of PNs and entraining them during network oscillations in the β-γ-frequencies range (Buzsaki and Wang, 2012;Cardin et al., 2009;Sohal et al., 2009). Therefore, we hypothesized that changes of perisomatic inhibitory strength might affect the temporal association of PN firing with ongoing γ-activity. Optogenetic activation of ChR2-expressing layer 2/3 PNs (Figure 4 – figure supplement 1) with ramps of blue light evokes robust γ-oscillations that depend on both GABAergic and glutamatergic synaptic transmission (Adesnik and Scanziani, 2010;Pouille et al., 2009;Shao et al., 2013). Importantly, photo-induced γ-activity is faithfully transmitted vertically to layer 5 neurons belonging to the same cortical column (Adesnik and Scanziani, 2010).

We induced oscillatory activity by photostimulation of ChR2+ layer 2/3 PNs, while simultaneously recording IPSCs and EPSCs in voltage-clamp in ChR2-negative PNs in layers 2/3 and 5, respectively (average amplitude of IPSCs: 703.9 ± 82.45 pA; average amplitude of EPSCs: 266.8 ± 56.87 pA; n = 15, Figure 4A and B). Photostimulation induced robust oscillations in the γ-frequency range involving both IPSCs and EPSCs, in layers 2/3 and 5 PNs, respectively (frequency IPSCs = 26.04 ± 0.78 Hz, frequency EPSCs = 27.26 ± 1.37 Hz n = 15, p = 0.45, Wilcoxon matched-pairs signed rank test, Figure 4C). We then recorded layer 5 PNs in current-clamp mode, in order to analyze how their spikes were temporally associated to ongoing IPSC rhythmic activity, recorded in voltage-clamp in layer 2/3 PNS (Figure 4D and E). During baseline, layer 5 PNs were tuned to γ-oscillations and discharged on average ∼2 ms before the peak of the γ-cycle (spike time to oscillations peak during baseline: −2.44 ± 0.44 ms, n = 9, Figure 4E and F). Remarkably, after inducing LTPi-FFI with bursts of APs, layer 5 PN firing was significantly anticipated (spike time to oscillations after AP bursts: −4.22 ± 0.52 ms, n = 9, p = 0.0313, Wilcoxon matched-pairs signed rank test, Figure 4E and F). The observed change in spike timing was not due to changes in firing rate nor resting membrane potential (Figure 4G and H, p = 0.08 for Bsl vs LTPi firing rate; p = 0.49 for Bsl vs LTPi membrane potential, paired *t* test). Interestingly, in cells, in which LTPi-FFI could not be induced, we did not observe a change in timing association (Baseline time to oscillations peak, −5.33 ± 1.12 ms; after bursts, −5.67 ± 1.2 ms, n = 6, p = 0.36, paired *t* test, Figure 4F). These results indicate that potentiation of feed-forward perisomatic inhibition alters the temporal association of layer 5 PNs during γ-oscillations.

### Bursts of APs decrease the firing rate in a subset of layer 5 PNs in vivo

We have shown that LTPi-inducing bursts decrease output spike probability and frequency of layer 5 PNs (Lourenço et al., 2014) (Figures 2 and 3). We wanted to test if such modulation occurred *in vivo*. We performed whole-cell recordings from layer 5 PNs in the barrel cortex of anesthetized mice (see Materials and methods) during spontaneous activity and in the presence of sensory stimulations, induced by air puffs to the contralateral whisker pads. Location and PN identity was confirmed by anatomy in some experiments (Figure 5 - figure supplement 1). PNs could be separated into two populations, according to their change in firing rate following AP bursts (5APs at 100Hz, repeated 15 times every 10 seconds). Type A cells displayed increased firing rate upon LTPi–inducing bursts during spontaneous activity and in the presence of sensory stimulations (1.37 ± 0.28 vs. 2.43 ± 0.52 Hz, pre-bursts vs. after bursts, respectively; n = 10, p = 0.002; Wilcoxon matched-pairs signed rank test; Figure 5A,B; Figure 5 - figure supplement 2). Conversely, type B cells exhibited a marked decrease in firing rate (2.4 ± 0.56 vs 1.04 ± 0.26 Hz, pre-bursts vs. after bursts, respectively; n = 16, p <0.0001Wilcoxon matched-pairs signed rank test; Figure 5C,D; Figure 5 - figure supplement 2). Importantly, these changes were not associated with significant variations in membrane potential (Type A: −57.9 ± 2.8 vs. −55.4 ± 2.9 mV, pre-bursts vs. after bursts; n = 10, p = 0.08, Wilcoxon matched-pairs signed rank test; Type B: −58.09 ± 2.9 vs. −56.31 ± 2.0 mV, pre-bursts vs. after bursts; n = 16, p = 0.25, Wilcoxon matched-pairs signed rank test). Type A and B accounted for 38.5 and 61.5% of recorded PNs.

**Figure 5.**
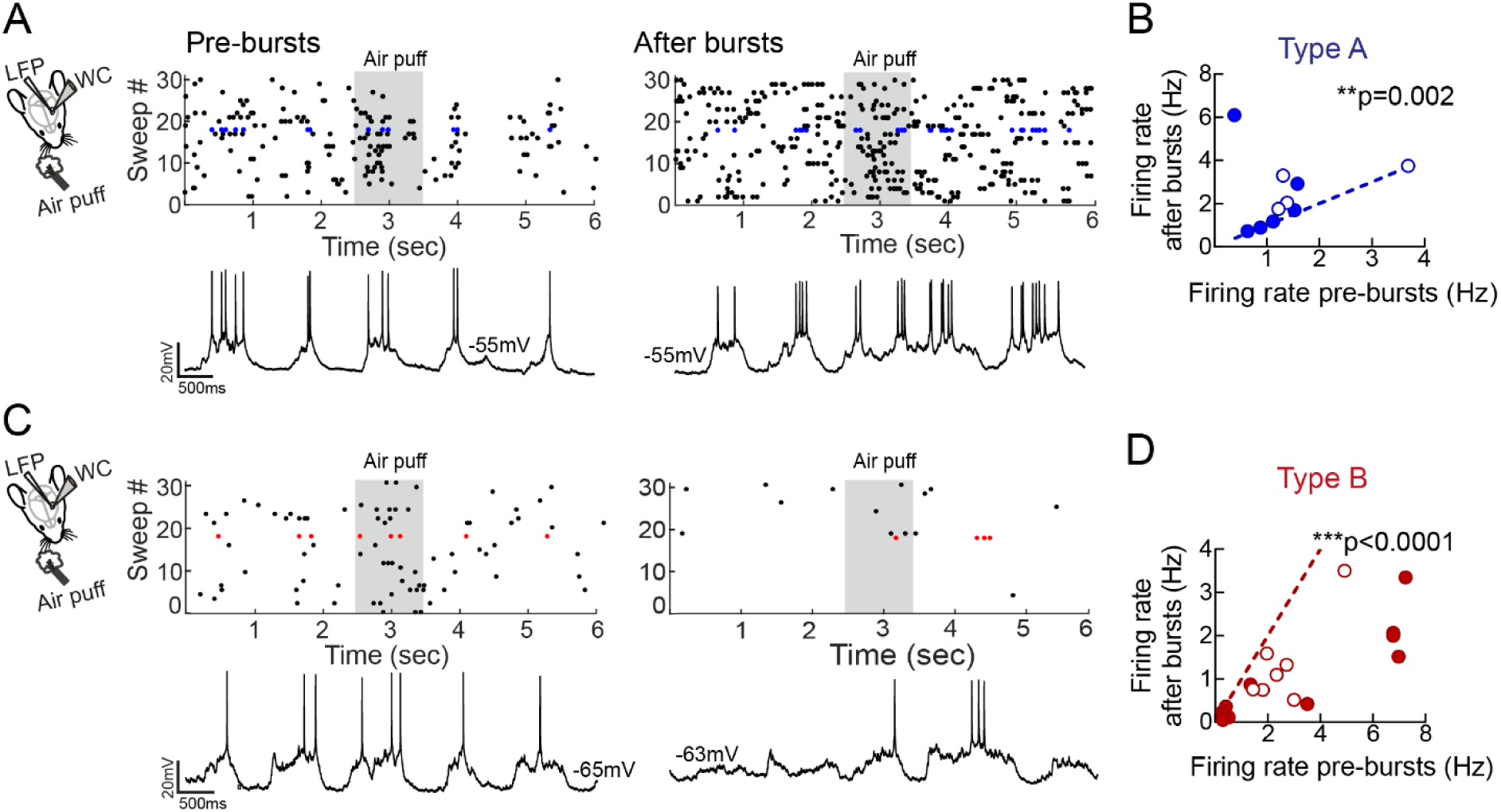
Bursts of APs decrease the firing rate in a subset of layer 5 PNs in vivo. (**A**) Left panel, Scheme of the recording configuration: LFP and whole-cell (WC) recordings of layer 5 PN in S1, during spontaneous activity and upon air-puff stimulation. Right: representative experiment of a type A PN (top: raster plot; bottom: representative current-clamp trace). Air puff stimulation are indicated as a gray rectangle, during a pre- and post-bursts period. (**B**) Plot of mean firing rate from individual PNs before (x-axes) vs. 5-10 min after postsynaptic bursts (y-axes). Type A layer 5 PNs displayed a small albeit significant increase in firing rate after bursts. Dotted line indicates unitary values (no change). Blue-filled symbols refer to PNs receiving air-puff stimulation; open circles refer to PNs, which did not receive air-puff stimulations. Note that these are different cells, from different recordings.**p<0.01, Wilcoxon matched-pairs signed rank test. (**C**-**D**) same as in (**A**-**B**), but for type B cells, in which postsynaptic bursts induced a decrease in firing rate in both conditions. ***p<0.0001, Wilcoxon matched-pairs signed rank test.

These results indicate that, in vivo, a prominent fraction of PNs respond to repetitive AP burst firing with decreased firing rate, possibly through potentiation of perisomatic inhibition, similarly to our slice results.

### Reduced firing rate is associated to increased tuning of PN spiking activity with γ-oscillations in vivo

We found that LTPi is associated to altered temporal association of PN firing to photo-induced γ-oscillations in acute cortical slices (Figure 4). We therefore examined whether postsynaptic bursting activity changes the temporal association of layer 5 PN firing with rhythmic activity in vivo. Previous evidence indicate that cortical PNs are poorly coupled to spontaneous and sensory-evoked γ-activity (Perrenoud et al., 2016). Could increases of perisomatic inhibition facilitate the tuning of PN spiking with ongoing network oscillations? In vivo whole-cell recordings of layer 5 PNs were coupled to local field potential (LFP) recordings obtained with a separate electrode (Figure 6 – figure supplement 1; see Materials and methods). We analyzed the relationship of spike probability of layer 5 PNs to γ-activity embedded in the LFP before and after bursting in the presence and absence of sensory stimulations. As previously reported in the visual cortex (Perrenoud et al., 2016), PN spiking activity was poorly tuned both in the presence and absence of whisker stimulations (Figure 6 and Figure 6 - figure supplement 2). On average, spike distributions did not reveal a significant phase preference under both conditions (Figure 6 and Figure 6 - figure supplement 2). However, in 8 out of 11 cells recorded during spontaneous activity (n = 10 mice) and in 11 out of 15 cells recorded during air puff stimulations (n = 13 mice), we observed a slight but significant increase of spike tuning with γ-activity. This was indicated by nonrandom spike-phase distributions crossing the Monte Carlo simulation threshold (see Materials and methods; Figure 6 and Figure 6 - figure supplement 2).

**Figure 6.**
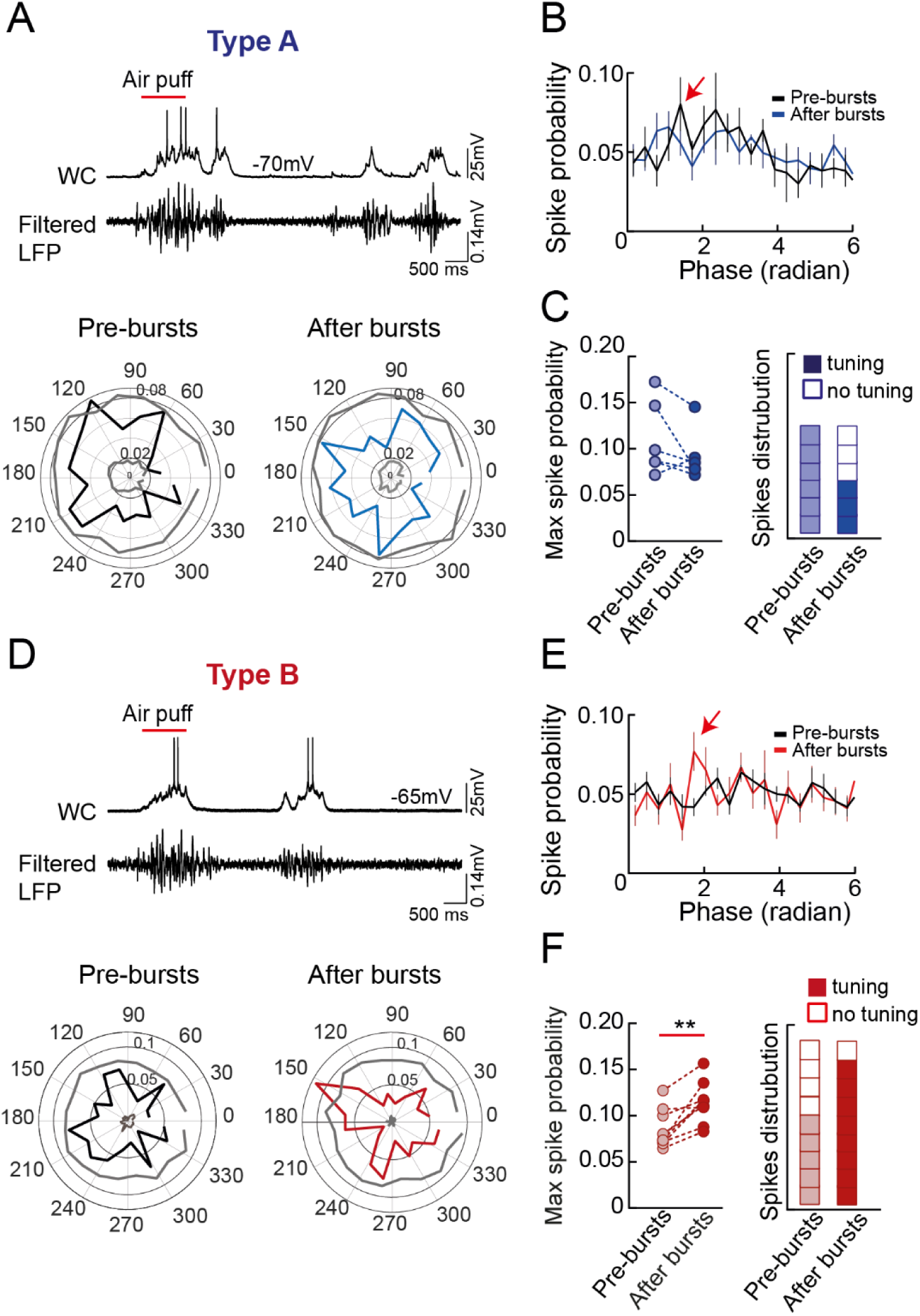
Reduced firing rate is associated to increased tuning of PN spiking activity with γ-oscillations in vivo. (**A**) Top, representative current-clamp traces showing firing activity in a Type-A layer 5 PN (top trace), and filtered LFP (between 20 and 100 Hz; bottom trace). The red bar indicates the air-puff stimulation. Bottom: representative polar plots displaying the distribution of APs with γ-phases in the same Type A neuron before (left, pre-bursts, black trace) and after bursts (right, post-bursts, blue trace). The gray lines correspond to the confidence intervals (set at 1.5 SD threshold) estimated by Monte Carlo methods from 100 independent surrogate data set. (**B**) Population spike distribution across the phase before (black trace) and after bursts (blue trace). Please note the point of maximum (max) spike probability (red arrow). (**C**) Left: summary plot illustrating the max spike probability extracted from each cell before (light blue circles) and after bursts (dark blue circles). Right: plot of the number of cells before (light blue) and after bursts (dark blue) whose spike circular distributions crossed the confidence intervals as shown in A, bottom. White boxes represent non-tuned PNs. (**D**-**F**) same as in (**A**-**C**) but for type B cells. Please note that in these PNs postsynaptic bursts induced a marked increase in max spike probability. In (**B**) and (**E**), the error bars indicate SEM. **p<0.01, Wilcoxon matched-pairs signed rank test.

Interestingly, in sensory-evoked conditions, whereas type A cells did not exhibit any significant change in max spike probability (Figure 6A-C), type B displayed a significant increase in max spike probability after bursts (Figure 6D-F; Type A: 0.11 ± 0.016 vs. 0.092 ± 0.011, baseline vs. after bursts; n = 6, p = 0.22, Wilcoxon matched-pairs signed rank test; type B: 0.084 ± 0.007 vs. 0.113 ± 0.007, baseline vs. after bursts; n = 9, p = 0.0039, Wilcoxon matched-pairs signed rank test). Accordingly, after AP bursts, the fraction of type B neurons tuned to γ-activity increased (5 out of 9 vs. 8 out of 9, baseline vs. after bursts, respectively). This was reflected also by an increased delta pairwise phase consistency (PPC), which is a measure of phase locking, independent from spike rate (Perrenoud et al., 2016;Veit et al., 2017) (Figure 6 – Figure 6 supplement 3).

These results indicate that type-B layer 5 PNs increase their tuning to γ-oscillations both during spontaneous and sensory-evoked activity. This suggests that likely potentiation of inhibitory transmission promotes orchestrated activity of single PNs in vivo.

### A computational model reveals two separate effects of LTPI-FFI

Our *in vitro* and *in vivo* results (Figures 4 and 6) suggest that LTPi alters the temporal association of PN firing during γ-oscillations. This prompts the question whether this is due to the actual strength of PN perisomatic inhibition, which sets the actual E/I level. To address this question, we examined a computational model of the feed-forward inhibition circuit between layers 2/3 and 5 (Figure 7A). In this model, a layer 5 PN was represented by an integrate-and-fire neuron, which was driven by layer 2/3 oscillatory activity (30 Hz) through a composite PSC (Figure 7B). As determined experimentally in response to brief layer 2/3 optical stimulations (Figure 1A-C) or synaptic recordings (Lefort et al., 2009), the composite PSC consisted of an early excitatory component, followed after a short delay by an inhibitory component corresponding to FFI (see Materials and Methods). The amplitude of the inhibitory component was systematically varied to investigate the effects of LTPi-FFI in the model. Additional inputs to the layer 5 PN were modeled as background noise.

**Figure 7.**
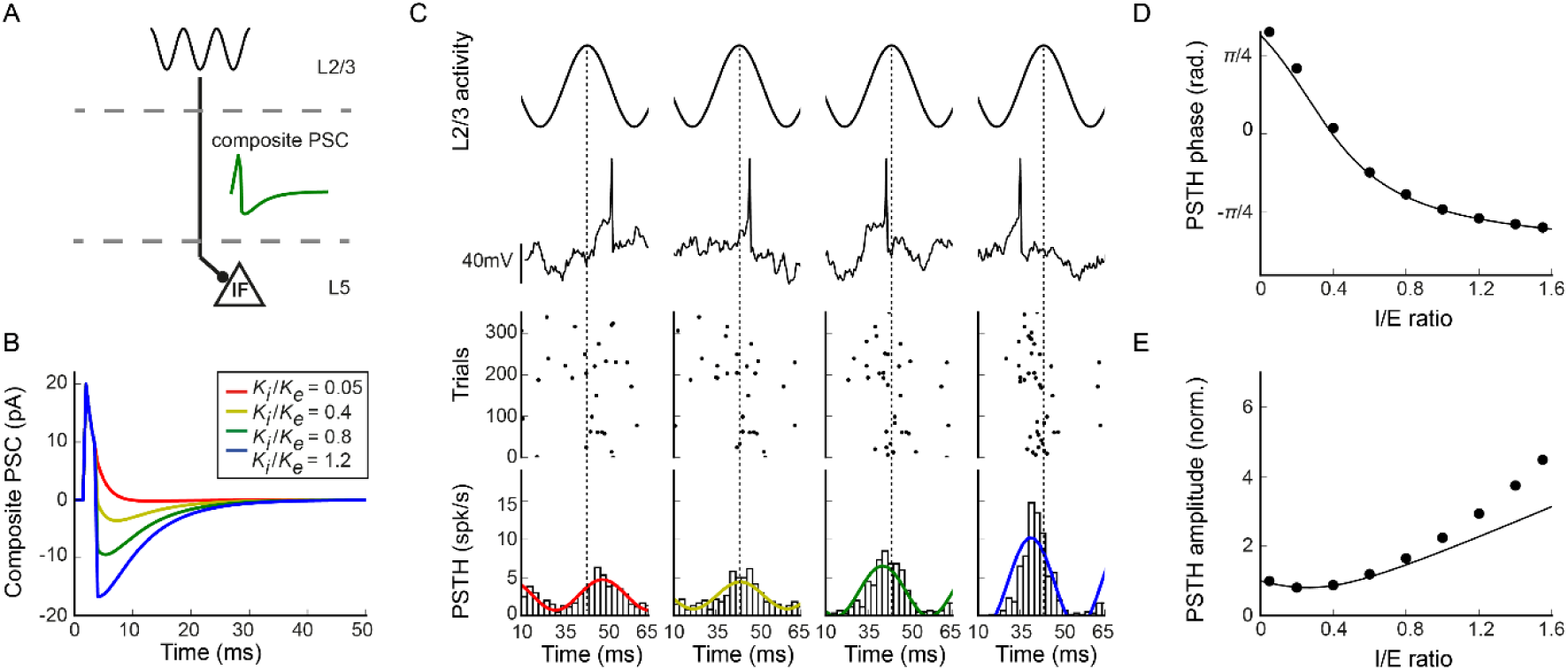
A computational model reveals two separate effects of LTPI-FFI. (**A**) Schematic of the model. (**B**)Composite postsynaptic current received by the model layer 5 PN in response to a brief layer 2/3 activation, for four increasing values of the inhibitory strength. (**C**) Activity in the model layer 5 PN in response to oscillatory layer 2/3 inputs, for four increasing values of the inhibitory strength (left to right). From top to bottom: layer 2/3oscillation; illustration of membrane potential traces in the model layer 5 PN; rastergram of model layer 5 PN action potentials over 300 repeats of the input; histogram of the average activity computed over 3000 repeats of the input. (**D-E**) Phase and amplitude of the average layer 5PN activity as function of the inhibition/excitation ratio. Dots: results of cosine fits to simulation data (illustrated in the bottom panels in C); continuous line: theoretical prediction based on the shape of the composite PSC (see Methods). In D, the phase is determined with respect to layer 2/3 input. In E, the amplitude was normalized to 1 for a inhibition/excitation ration of 0.05. Note that the theoretical prediction is based on a linear approximation. This approximation fails at large values of inhibition, as seen in the deviations between predicted and measured amplitude (panel **E**). Non-linear effects lead to an increase in PSTH oscillation amplitude, and therefore firing precision, with respect to the linear prediction.

Layer 2/3 oscillatory inputs filtered through the composite PSC led to a temporal modulation of the activity in the model layer 5 PN (Figure 7C). To determine the temporal relationship between layer 5 and layer 2/3 activity, we computed the histograms of temporal activity in layer 5, and extracted its phase and amplitude by fitting cosine functions (Figure 7C). The phase of the histogram quantifies the average timing of layer 5 activity with respect to layer 2/3 oscillations, while the amplitude of the modulation represents the precision of layer 5 spikes. A mathematical analysis of the circuit dynamics predicted that the phase and amplitude of layer 5 PN activity are determined by the phase and amplitude of the Fourier transform of the composite PSC, evaluated at the frequency of 30 Hz that corresponds to layer 2/3 oscillatory inputs (Figure 7B, see Materials and Methods). A comparison with numerical simulations confirmed this prediction (Figure 7D and E).

The circuit model revealed that increasing the amplitude of the inhibitory component of the composite PSC led to two separate effects. The first effect is that increasing inhibition at moderate strengths advances the average timing of layer 5 activity with respect to layer 2/3 oscillations, but leaves the precision of the activity essentially unchanged. This timing shift is consistent with the experimental measurements reported in Figure 4 (note that in Figure 4 the absolute timing was quantified with respect to layer 2/3 IPSCs, while in the model the timing is quantified with respect to layer 2/3 average firing activity). The model shows that the range of possible timing shift is limited, as the phase shift saturates as inhibition is increased. Interestingly, however, large increases in inhibition therefore lead to a second, separate effect, in which the timing of layer 5 activity does not change anymore, but its precision increases with stronger inhibition, as shown previously (Lourenço et al., 2014). Accordingly, this effect is consistent with increased tuning in vivo shown in Figure 6.

## Discussion

In this study, we provide evidence that bursting activity of layer 5 PNs can potentiate incoming feed-forward inhibition recruited by a descending excitatory pathway, which is one of the most prominent in the neocortex. This plasticity likely originated from PV basket cells, strongly modulated PN gain, affected coordinated activity across cortical layers and altered the temporal association of PNs with γ-oscillations both *in vitro* and *in vivo*.

In principle, feed-forward inhibition can result from long-range recruitment of different interneuron subclasses (Pluta et al., 2019). Yet, potentiation of FFI likely involves perisomatic inhibition originating almost exclusively from PV basket cells. Indeed, here we provide evidence that optogenetic activation of layer 2/3 PNs efficiently recruits layer 5 PV basket cells (Figure 1 – supplement figure 1). Moreover, LTPi-FFI was completely abolished by the canonical NO receptor inhibitor ODQ, which we previously showed to be selective for PV cell-mediated LTPi, and did not involve dendrite-targeting somatostatin (SST)-positive interneurons (Lourenço et al., 2014). Moreover, SST interneurons are recruited more efficiently in response to prolonged trains, due to strongly facilitating glutamatergic synapses (Kapfer et al., 2007;Silberberg and Markram, 2007;Wang et al., 2004). Although we cannot exclude that other interneuron types might be involved in LTPi-FFI, we believe that this plasticity relies on the strength of PV cell synapses. Accordingly, PV basket cells were demonstrated to be the most prominent, if not exclusive, providers of perisomatic inhibition onto large deep layer PNs of somatosensory cortex (Bodor et al., 2005).

Importantly, postsynaptic bursting activity potentiated feed-forward inhibition strongly, without affecting the strength of the glutamatergic component of the composite synaptic response. Our finding that PNs can unlock the E/I ratio is in line with our previous report on selective potentiation of GABAergic synapses from PV cells, through retrograde nitric oxide signaling (Lourenço et al., 2014). This is the first time, to our knowledge, that postsynaptic firing activity alone can effectively alter the E/I ratio of a prominent input pathway, impinging layer 5 PNs.

A tight balance between excitation and inhibition is believed to guarantee the proper functioning of neural circuits (Isaacson and Scanziani, 2011;Marin, 2012), particularly in primary sensory cortical areas, such as the somatosensory cortex (Higley and Contreras, 2006;Okun and Lampl, 2008), mature auditory cortex (Froemke et al., 2007;Wehr and Zador, 2003) and visual cortex (Ferster, 1986;Hensch and Fagiolini, 2005). However, the loosening of the E/I lock is necessary for sensory processing and the refinement of sensory maps (Froemke et al., 2007). Moreover, it has been shown that E/I ratio is different across individual cortical principal cells, depending on the specific layer (Adesnik and Scanziani, 2010), and the intrinsic activity state of each PN (Xue et al., 2014), demonstrating that the tight lock of E/I ratio can be disrupted by perturbing pyramidal cell activity. Therefore, plasticity of GABAergic perisomatic FFI can be a relevant mechanism to tune PN sensitivity to process sensory information within the cortical column.

We found that the selective increase of perisomatic inhibitory strength onto a PN, without altering its excitatory input, has a major effect on input-output (I-O) relationship. Indeed, LTPi-FFI results in a strong divisive output modulation of the I-O curve, as it significantly changed its slope, without altering the rheobase. From a computational perspective, this means that, for rate-coded neuronal signaling, the modulation operated by LTPi-FFI results in a change of PN gain that affects the dynamic range of its response, likely preventing saturation of firing (Silver, 2010). However, whether inhibition play an additive or multiplicative role has been debated, depending on the location (perisomatic vs. dendritic) and strength of the inhibitory response, as well as on the morphological complexity of the postsynaptic neuron (Lovett-Barron et al., 2012;Silver, 2010). In particular, perisomatic PV cell-mediated modulation of orientation tuning of visual cortical PNs was shown to have both an additive and multiplicative effect, with a strong modulation of PN gain during sensory processing (Atallah et al., 2012;Lee et al., 2012;Wilson et al., 2012). Here we show that cell-autonomous potentiation of perisomatic inhibition has a strong effect on PN gain. This was not due to changes of intrinsic excitability (e.g. changes in membrane potential or resistance), but to selective increase of feed-forward inhibition. Therefore, by inducing retrograde potentiation of perisomatic inhibition, burst firing of PNs altered their functional network connectivity, and thus the dynamic pattern of their activation, through a divisive modulation of their gain.

Cortical PNs are known to respond to sensory stimuli using a sparse population coding (Petersen and Crochet, 2013). In this regime, neurons act as coincidence detectors of temporally correlated input: changes of the gain of the I-O curve of PNs might shape the time window in which inputs can be integrated to generate a spike (Lourenço et al., 2014;Pouille and Scanziani, 2001). It is therefore likely that burst firing-induced potentiation of inhibition controls the temporal properties of signals propagating through the network.

The dynamic modulation of PN firing, induced by plasticity of feed-forward inhibition had a powerful effect on the columnar integration of activity across cortical layers. Indeed, activation of layer 2/3 PNs was shown to increase the firing of PNs in layer 5 of the same column (Adesnik and Scanziani, 2010; but see Pluta et al., 2019). LTP of feed-forward inhibition strongly diminished (in fact, on average, it abolished) the layer 2/3-dependent activation of layer 5 PNs, thereby affecting the flow of neuronal communication across cortical layers.

Here we show that layer 5 PNs are tuned to γ-activity, optogenetically generated in layer 2/3. Indeed, PNs spiked at a relatively precise time in reference to photo-activated network rhythmic activity, but the induction of long-term plasticity of feed-forward perisomatic inhibition shifted the temporal association of layer 5 PN spikes with γ-oscillations. This effect was due to increased inhibition, as glutamatergic neurotransmission, overall PN firing rate and passive properties were not altered by postsynaptic burst firing.

Photo-activated γ-oscillations represents a very useful tool to dissect the cellular and synaptic mechanisms underlying network synchronization (Adesnik and Scanziani, 2010;Quiquempoix et al., 2018). However, this approach suffers from some limitations, namely hyper-synchronous activity induced by simultaneous prolonged activation of a large number of neurons in a reduced preparation. We therefore investigated whether LTPi-inducing trains could affect spike timing relative to endogenous γ-activity in a more intact preparation. We found that our *in vitro* and *in vivo* results converge: burst firing induced a decrease in firing rate in a prominent fraction of layer 5 PNs, and improved tuning during both spontaneous and sensory-evoked γ-activity. The modulation of spike association to network oscillations both *in vitro* and *in vivo* that we show here is in line with the known function of perisomatic inhibition (particularly from PV basket cells) to entrain PNs during γ-activity.

Moreover, both *in vitro* and *in vivo*, we found a similar low percentage of neurons, which did not display LTPi and did not exhibit decreased firing rates, respectively. We termed these neurons Type A in our *in vivo* recordings, and we speculate that they correspond to the fraction of PNs not expressing LTPi *in vitro*. In these PNs, coupling with network activity was not affected by burst firing. It is currently unclear whether these PNs belong to different cell types. Indeed, it has been demonstrated that thick-tufted large cortico-fugal PNs of the prefrontal cortex are preferentially innervated by PV cells, as compared to more slender, thin-tufted cortico-cortical PNs (Allene et al., 2015;Lee et al., 2014). Alternatively, in this minority of PNs, lack of LTPi expression could be due to an already potentiated (and hence saturated) inhibition. This could explain the consistent negative shift of their spike times in vitro, as compared to PNs, which did undergo LTPi. Moreover, bi-directional, activity-dependent changes of firing were reported in layer 5 PNs of rat neocortex (Mahon and Charpier, 2012).

Importantly, our computational model revealed that increasing inhibition in the feed-forward circuit induced two separate effects for different levels of the E/I ratio. If excitation dominates, increasing inhibition modulates the timing of layer 5 spikes, but not their precision with respect to layer 2-3 oscillations. This is indeed what we found during photo-activated γ-oscillations in neocortical slices. If, however, inhibition becomes stronger than excitation, further increases in inhibition improve the precision of layer 5 spikes (as found in (Lourenço et al., 2014)), but do not additionally affect their timing. This could explain the improvement in tuning that we detect *in vivo*, where assessing the actual E/I ratio is technically challenging and prone to strong biases. Modifying inhibition strength within different ranges may therefore lead to different functional consequences. Note that the strength of inhibitory connections is known to control coherence in recurrent networks (Bartos et al., 2002;Brunel and Hakim, 1999;Brunel and Wang, 2003). Here in contrast we demonstrated its specific role for the temporal organization of activity in FFI circuits.

Therefore, plasticity of feed-forward perisomatic inhibition, by governing spike-time association of single PNs with ongoing γ-activity, might be responsible for shifting their participation to distinct cell assemblies, thereby reconfiguring the local network (Mongillo et al., 2018).

In conclusion, LTPi of feed-forward inhibition can be a simple mechanism modulating the functional connectivity of single PNs, largely influencing cortical networks and subnetworks. LTPi of PV cell-mediated feed-forward inhibition in sensory cortices could be a fundamental dynamic property of cortical networks, providing the basis of diverse cognitive functions, such as sensory perception and attention.

## Materials and methods

### Animals

Experimental procedures followed national and European (2010/63/EU) guidelines, and have been approved by the authors’ institutional review boards and national authorities. All efforts were made to minimize suffering and reduce the number of animals. Experiments were performed on C57BL/6 wild-type mice (Janvier Labs, France).

### In utero electroporation

Timed-pregnant C57BL/6 wild-type female mice (15.5 days postcoitum) were anaesthetized with 1-2% isoflurane. The abdomen was cleaned with 70% ethanol and swabbed with betadine. Buprenorphine (0.05 mg/kg) was administered subcutaneously for preoperative analgesia and local anesthetic bupivacaine (2.5mg/kg) was injected between the skin and the abdomen 5 min before incision. A midline ventral laparotomy (∼2 cm) was performed, and the uterus gently exposed and moistened with PBS, pre-warmed at 37 °C. Using glass beveled capillaries, DNA plasmids mixed in saline (PBS) solution and 0.025% Fast Green (Sigma) were injected through the uterine wall into the lateral ventricle of each embryo. Embryos were injected with pCAG-mRFP (0.8 μg/μl) (Manent et al., 2009) (Addgene #28311) plasmid DNA mixed with either pCAG-ChR2-Venus (Petreanu et al., 2007) (Addgene #15753) or pCAG-ChETA-EYFP (1.5μg/μL). ChETA-EYFP was subcloned from p-Lenti-CaMKIIa-ChETA-EYFP (Gunaydin et al., 2010) (Addgene #26967) into the backbone of pCAG-ChR2. After each injection, the embryos were moistened with PBS. DNA was electroporated via 5 square electrical pulses of 40 V amplitude and 50 ms duration through forceps-type circular electrodes positioned at 0° angle with respect to the rostral-caudal axis of the head of the embryos. After electroporation, the uterus was placed back into the peritoneal cavity and moistened with PBS. The abdomen and skin were then sutured and the latter cleaned with betadine. The procedure typically lasted maximum 40 min starting from anesthesia induction. Pups were born by natural birth and placed with a Swiss foster mother (timed-pregnant at the same time as C57BL/6 females) after they were screened for location and strength of transfection by trans-cranial epifluorescence under a fluorescence stereoscope.

### *In Vitro* Slice Preparation and Electrophysiology

Coronal slices (400-μm-thick) from somatosensory cortex were obtained from 15- to 28-day-old mice. Animals were deeply anesthetized with isofluorane and decapitated. Brains were quickly removed and immerse in “cutting” solution (4°C) containing the following (in mM): 87 NaCl, 25 NaHCO_3_, 2.5 KCl, 1.25 NaH_2_PO_4_, 7.5 MgSO_4_, 0.5 CaCl_2_, 2 pyruvic acid, 3 myo-inositol, 0.4 ascorbic acid, 25 glucose and 70 sucrose (equilibrated with 95% O_2_ / 5% CO_2_). Slices were cut with a vibratome (Leica) in cutting solution and then incubated in oxygenated artificial cerebrospinal fluid (ASCF) containing the following (in mM): 125 NaCl, 3 KCl, 2.5 CaCl_2_, 1.3 MgSO_4_, 1.25 mM NaH_2_PO_4_, 26 mM NaHCO_3_, 0.4 ascorbic acid and 16 mM glucose (pH 7.4), initially at 34°C for 30 min, and subsequently at room temperature, before being transfer to the recording chamber. Recordings were obtained at 30°C. Synaptic events were recorded in whole-cell, voltage- or current-clamp mode from layer 2/3 and layer 5 PNs of mouse primary barrel somatosensory cortex visually identified using infrared video microscopy (Lourenço et al., 2014). For voltage clamp experiments of layer 2/3 PNs, electrodes (with a tip resistance of 2-4 MΩ) were filled with a cesium-based internal solution (in mM): 120 CsMeSO_4_, 8 CsCl, 10 HEPES, 10 BAPTA, 4 NaCl, 2 CaCl_2_, 4 Mg-ATP, 0.3 Na-GTP, 4 phosphocreatine di(tris), 0.5 QX-314-Cl; pH adjusted to 7.2 with CsOH; 280-300 mOsm. Under these recording conditions, activation of GABA_A_ receptors resulted in outward currents at a holding potential (*V*_h_) of +10 mV. In current clamp experiments electrodes were filled with a potassium-based intracellular solution containing (in mM): 130 K-gluconate, 5 KCl, 10 HEPES, 0.2 EGTA, 2 MgCl_2_, 4 Mg-ATP, 0.3 Na-GTP, 4 phosphocreatine di(tris); pH adjusted to 7.2 with KOH; 280-300 mOsm. In current-clamp mode cells were recorded at their resting membrane potential unless for Figure 4 where occasionally depolarization (max. current injection 30pA) was required in order to induce firing of layer 5 PN upon layer 2/3 light activation. In voltage-clamp experiments, access resistance was on average <20 MΩ and monitored throughout the experiment. Recordings were discarded from analysis if the resistance changed by >20% over the course of the experiment. In current-clamp experiments, input resistance was monitored with small current steps (−30 pA for 600 ms) and cells were excluded if it changed by >25%. ODQ was obtain from R&D Systems Europe.

## Data analysis

Signals were amplified, using a Multiclamp 700B patch-clamp amplifier (Axon Instruments, Foster City, California, United States), sampled at 50 kHz and filtered at 4 or 10 kHz for voltage and current-clamp mode, respectively. Data were analyzed using pClamp (Axon Instruments), IGOR PRO 5.0, (Wavemetrics), MATLAB (MathWorks) and GraphPad Prism software.

### IPSCs during rhythmic activity *in vitro*

Custom written software (Detector, courtesy J. R. Huguenard, Stanford University) was used for analyzing GABAergic events, as previously described (Manseau et al., 2010;Ulrich and Huguenard, 1996). Briefly, individual events were detected with a threshold-triggered process from a differentiated copy of the real trace. For each cell, the detection criteria (threshold and duration of trigger for detection) were adjusted to ignore slow membrane fluctuations and electric noise while allowing maximal discrimination of IPSCs. Detection frames were regularly inspected visually to ensure that the detector was working properly.

### Spike probability of layer 5 PNs during rhythmic activity *in vitro*

Analysis of the temporal relationship between layer 5 PNs spikes and ongoing rhythmic activity was analyzed using custom written scripts in MATLAB (The MathWorks, Inc., Natick, Massachusetts, United States). Briefly, spikes were extracted using a threshold of −10 mV on the membrane potential trace, and the times of the action potential peaks were extracted after cubic spline data interpolation of the waveforms around the spikes. Next, IPSC peak positions from the detector (see above) were adjusted with a 50-samples smoothing of the current waveform, and delays between action potential timing and IPSC occurrence were computed. Finally, histograms between spike times and IPSC occurrences were generated using 2 ms binning and converted into probability distributions.

#### Photostimulation

ChR2 or ChETA activation was induced by light flashes on cortical slices, using a 20 mW LED (λ = 470 nm, Cairn research, UK) collimated and coupled to the epifluorescence path of a Zeiss AxioExaminer microscope, using a 40X water immersion (N.A. 1) lens. In order to trigger robust oscillations (Figure 4), light ramps had a duration of 1–2 s, started at zero intensity and reached a final intensity of 9 mW/mm^2^. The stimulus intensity was adapted to each slice, depending on the opsin expression and it was repeated with a frequency of 0.025 Hz. ChR2 activation was also obtained by brief square light pulses (ranging between 0.5 and 1 ms) evoking postsynaptic potentials in layer 5 PNs (Figure 1).

#### Immunofluorescence

In order to check proper electroporation of both plasmids in the somatosensory cortex, in some cases (Figure 1A), slices used for electrophysiology experiments were fixed overnight in 4% paraformaldehyde in phosphate buffer (PB, pH 7.4) at 4°C. Slices were then rinsed three times at room temperature (10 min each time) in PB and were then rinsed three times in PB (10 min each) at room temperature and coverslipped in mounting medium. Immunofluorescence was then observed with an ApoTome.2 microscope (Zeiss) and images were acquired using a 10x objective.

#### Preparation for *in vivo* electrophysiology

Two- to three-week-old naive C57BL/6 or Scnn1a x tdTomato mice of both sexes were anesthetized via intraperitoneal (i.p.) injection with 15% urethane (1.5 g/kg in sodium lactate ringer solution) and placed on a stereotaxic apparatus. The body temperature was constantly monitored and kept at 37 °C with a heating blanket. Eye ointment was applied to prevent dehydration. To ensure a deep and constant level of anesthesia, vibrissae movement, eyelid reflex, response to tail, and toe pinching were visually controlled before and during the surgery. Subcutaneous injections of atropine (0.07 mg/kg) and dexamethasone (0.2mg/kg) were used to maintain clear airways and prevent edema, respectively. A mix of local lidocaine and bupivacaine injection was performed over the cranial area of interest and, after a few minutes, a longitudinal incision was performed to expose the skull. A stainless steel head post was sealed on to the mouse skull using dental acrylic cement. A small craniotomy (≥1 mm diameter) was made on the right hemisphere to target the primary somatosensory cortex according to stereotaxic coordinates at −1 from bregma: 3 lateral). And a second small craniotomy (approximately 200 μm far from the previous) was drilled for the separate entry of the LFP pipette. Dura was not removed. The exposed cortical surface was superfused with warm HEPES-buffered extracellular solution (in mM: 125 NaCl, 5 KCl, 10 glucose, 10 HEPES, 1.8 CaCl2 and 1 MgSO4 (pH 7.2)) in order to maintain ionic balance and prevent desiccation.

#### In Vivo LFP and whole-cell patch recording

Patch pipettes (5-7 MΩ) of 1.5 mm external diameter borosilicate glass (WPI) were pulled on a Narishige P100 Vertica Puller and filled with (in mM): 135 K-gluconate, 6 KCl, 10 HEPES, 1 EGTA, 4 MgATP, Na2ATP and 8 phosphocreatin, pH adjusted to 7.2 with KOH, 290-295 mOsm. Pipette capacitance was neutralized before break-in. Whole-cell patch-clamp recordings of L5 pyramidal neurons were performed following the standard techniques for blind patching (Margrie et al., 2002). High positive pressure was applied to the pipette to prevent tip occlusion. After breaking the meninges, the positive pressure was immediately reduced to prevent cortical damage. Once reached layer 5 depth,, the pipette was then advanced in 2-μm steps, and pipette resistance was monitored in the conventional voltage clamp configuration. When the pipette resistance suddenly increased, positive pressure was relieved to obtain a GΩ seal (small negative pressure can be applied to achieve seal formation). Seal resistances were always >1 GΩ. Recordings were made in current-clamp mode, and no holding current was applied. Typical recording durations were ∼15-20 min (which usually allowed 5 minutes recording of baseline and 5-10 minutes after burst trains). Air puff stimulation of the whisker pad was achieved by 1s-long pulses of compressed air delivered by a picospritzer unit via a 1 mm diameter plastic tube placed at ∼20 mm from the mouse snout. To record local field potential (LFP) patch pipette (1-2MΩ) were filled with HEPES-buffered extracellular solution and inserted in the cortex. Data were acquired at 50 kHz using a Multiclamp 700B Amplifier (Molecular Devices).

#### Biocytin filling

To certify that deep L5 PNs of S1 were being target, experiments were performed in Scnn1a x tdTomato mice allowing labelling of layer 4. Biocytin (Sigma) was added to the intracellular solution at a high concentration (0.5 g / 100 ml). Neurons were injected with large depolarizing currents in current clamp mode for fifteen times (100 ms, 1-2 nA, 1 Hz). Briefly, after *in vivo* experiments, mice were perfused and brain slices of 200 μm were performed as described above for *in vitro* electrophysiology. Slices were then fixed with 4% paraformaldehyde in phosphate buffer saline (PBS, Sigma) for at least 48 h. Following fixation, slices were incubated with the avidin-biotin complex (Vector Labs) and a high concentration of detergent (Triton-X100, 5%) for at least two days before staining with 3,3′Diaminobenzidine (DAB, AbCam) following (Jiang et al., 2015). Slices were then rinsed in PBS and coverslipped in mounting medium. Immunofluorescence and DAB staining were observed in a Micro Zeiss Routine microscope and images were acquired using a 10-40x objectives.

#### Analysis of the phase modulation of AP-LFP coupling

The analysis of the coupling between APs and local field potentials (LFP) was done in MATLAB (Mathworks, Natick MA, US), employing custom-written scripts. Briefly, AP occurrence times were extracted from the intracellular membrane potential, recorded and sampled at 50 kHz. This was based on a supervised peak-detection algorithm, using the findpeak() MATLAB function with a −15 mV detection threshold and a 4 ms dead-time. LFP were first down sampled to 1 KHz and then digitally filtered, between 20 and 200 Hz, by a 16^th^ order bandpass Butterworth filter. The waveform was then Hilbert-transformed and its instantaneous phase component unrolled in the range [0; 2*π*]. For each AP, the corresponding LFP phase was extracted as the corresponding value of the instantaneous phase component. A histogram was used as polar-plot or equivalently as a Cartesian-plot to display the distribution of AP-LFP phases. The confidence interval was estimated by Monte Carlo methods from 100 independent surrogate data set, obtained upon randomly jittering each AP times (i.e. Gaussian distributed, zero mean, 10 ms standard deviation) and again estimating the distribution of the corresponding LFP phases from the Hilbert instantaneous phase waveform.

#### Statistical analysis

Analysis of LTP was performed by comparing the mean amplitude of light evoked inhibitory postsynaptic currents in the 5 last minutes of the plasticity to the baseline period (Figure 1). Unless otherwise indicated, statistical comparisons were done between raw values.

Normality of the data was assessed (D’Agostino & Pearson omnibus normality test). Normal distributions were statistically compared using paired *t* test two-tailed and Wilcoxon matched-pairs signed rank test was used as a non-parametric test. Group data was analyzed using the Friedman test followed by Dunn’s multiple comparison test (e.g Figure 2D for different frequencies and Figure 3D). Differences were considered significant if p<0.05. Values are presented as mean ± SEM of *n* experiments.

Gain modulation (Figure 2F) was calculated from the average slope (F′) of the fits between 5% and 75% of its maximum value (Rothman et al., 2009). Only cells where the fit was possible were included in the calculation. Changes in gain (ΔGain) were computed as follows: 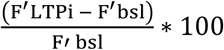. Additive offset shifts (ΔOffset) were defined as the difference between the half-maximum frequencies of the fits for the two conditions LTPi and baseline.

#### Computational model

Layer 5 PNs were modeled as integrate-and-fire neurons, with membrane potential dynamics given by

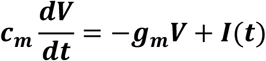

where the membrane potential ***V*** is determined with respect to the resting potential of the cell. The membrane capacitance was ***c***_***m***_=100 pF, and the membrane conductance ***g***=10 nS. An action potential was emitted when the membrane potential reached a threshold value V_T_=40 mV. The membrane potential was subsequently reset to a value V_R_ = 0 mV. The total input to the neuron was given by

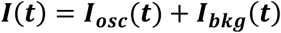

Here I_osc_ represents the oscillatory current received from layer 2/3 via feed-forward excitation and inhibition. It is therefore given by layer 2/3 oscillatory activity r_2/3_ (t),

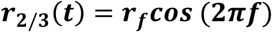

filtered by a composite post-synaptic current K(t), modeled as a difference of excitatory and inhibitory components

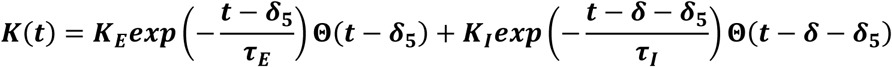

where ***τ***_***E***_= 2 ms and **τ**_***I***_= 7 ms are the timescales of excitation and inhibition, ***δ*** = 2 ms is the delay between excitatory and inhibitory inputs, ***δ***_**5**_ is an overall delay between layer 2/3 activity and layer 5 inputs, ***K***_***E***_ and ***K***_***I***_ represent the strengths of excitation and inhibition, and ***Θ***(***t***)is the Heaviside step function (0 for t<0, 1 for t>1).

The oscillatory input was therefore given by the convolution between K(t) and ***r***_**2**/**3**_(***t***):

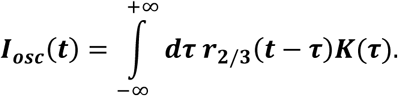

In the simulations, the amplitude of layer 2/3 oscillations was set to ***r***_***f***_=3 spk/s, the excitatory strength of the PSC was set to ***K***_***E***_=20 pA and ***K***_***I***_ was varied in the range from 0 to 32 pA, so that oscillations led to composite postsynaptic currents of about 100-350 pA.

Other inputs to the layer 5 PN were modeled as background noise

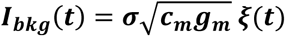

where ***ξ***(***t***) is Gaussian white noise of zero mean and unit variance, I_0_ is the mean input and ***σ*** is the background noise amplitude. In the simulations we used *σ*=22 *mv*

The oscillatory input from layer 2/3 entrains the activity of the layer 5 PN, which therefore acquires an oscillatory temporal structure. To determine the temporal relationship between layer 2/3 inputs and the output from layer 5PNs, we computed the trial-averaged firing rate ***r***_**5**_(***t***) (histograms in Figure 5 C bottom). If the oscillatory inputs are relatively weak, layer 5 PN activity can be approximated as

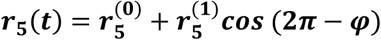

where the first term represents constant activity, and the second term describes an oscillation around that baseline. The phase ***φ*** quantifies the timing of the output with respect to the layer 2/3 oscillatory input, while the amplitude 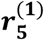 quantifies the precision of this timing – the larger 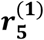, the more layer 5 PN action potentials are concentrated at the specific phase of the input given by ***φ*.**

Our aim was to understand how the shape of the composite post-synaptic current K(t) influences the timing and precision of the layer 5 PN output. Previous theoretical works [for a review, see (Brunel and Hakim, 2008)] showed that for weak inputs the phase and amplitude of the output can be decomposed into a synaptic and a neuronal contribution:

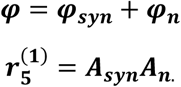

Here the neuronal contributions ***φ***_***n***_ and ***A***_***n***_ to the phase and amplitude depend on the background noise input and the specific neural model (Brunel et al., 2001;Fourcaud-Trocme et al., 2003). As we did not vary the background noise parameters, we treated them as arbitrary constants.

The synaptic phase lag ***φ***_***syn***_ and amplitude ***A***_***syn***_ are given by the phase and amplitude of the Fourier transform 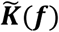 of the Fourier transform of the composite post-synaptic current K(t):

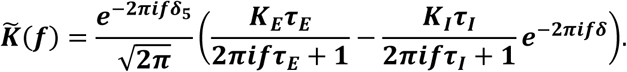

The phase and amplitude of 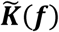 depend on the frequency of the oscillation (as well as on synaptic parameters). Since we were looking at the response to layer 2/3 inputs oscillating at 30Hz, 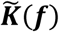 was evaluated at f=30 Hz, and a range of ***K***_***I***_/***K***_***E***_ values was used (see Figure 5).

The phase and the amplitude of the obtained values were then compared with direct fits of a cosine function to simulation results (Figure 5). Note that the above theoretical prediction for the phase and amplitude of layer 5 PN output rely on a linear approximation, expected to be accurate if the amplitude of the oscillations is not too strong. The comparison between the prediction and simulations indeed shows deviations from the predicted amplitude when inhibition becomes very strong (Figure 7 E).

## Acknowledgements

We thank Caroline Mailhes for initial help with *in utero* electroporation. We thank Michael Graupner and Christoph Schmidt-Hieber for the advices in establishing the in vivo whole-cell recording technique. This work was supported by European Research Council (ERC) under the European Community’s 7th Framework Programmme (FP7/2007-2013)/ERC grant agreement No 200808); “Investissements d’avenir” ANR-10-IAIHU-06; Agence Nationale de la Recherche (ANR-13-BSV4-0015-01; ANR-FRONTELS; ANR-NanoSynDiv; ANR-CaCoVi), Fondation Recherche Médicale (Equipe FRM DEQ20150331684), NARSAD independent investigator grant, and a grant from the Institut du Cerveau et de la Moelle épinière (Paris) (A.B.). SO was supported by the Programme Emergences of City of Paris, and the program “Investissements d’Avenir” launched by the French Government and implemented by the ANR, with the references ANR-10-LABX-0087 IEC and ANR-11-IDEX-0001-02 PSL Research University.

## Supplemental Data

**Figure 1-figure supplement 1:**
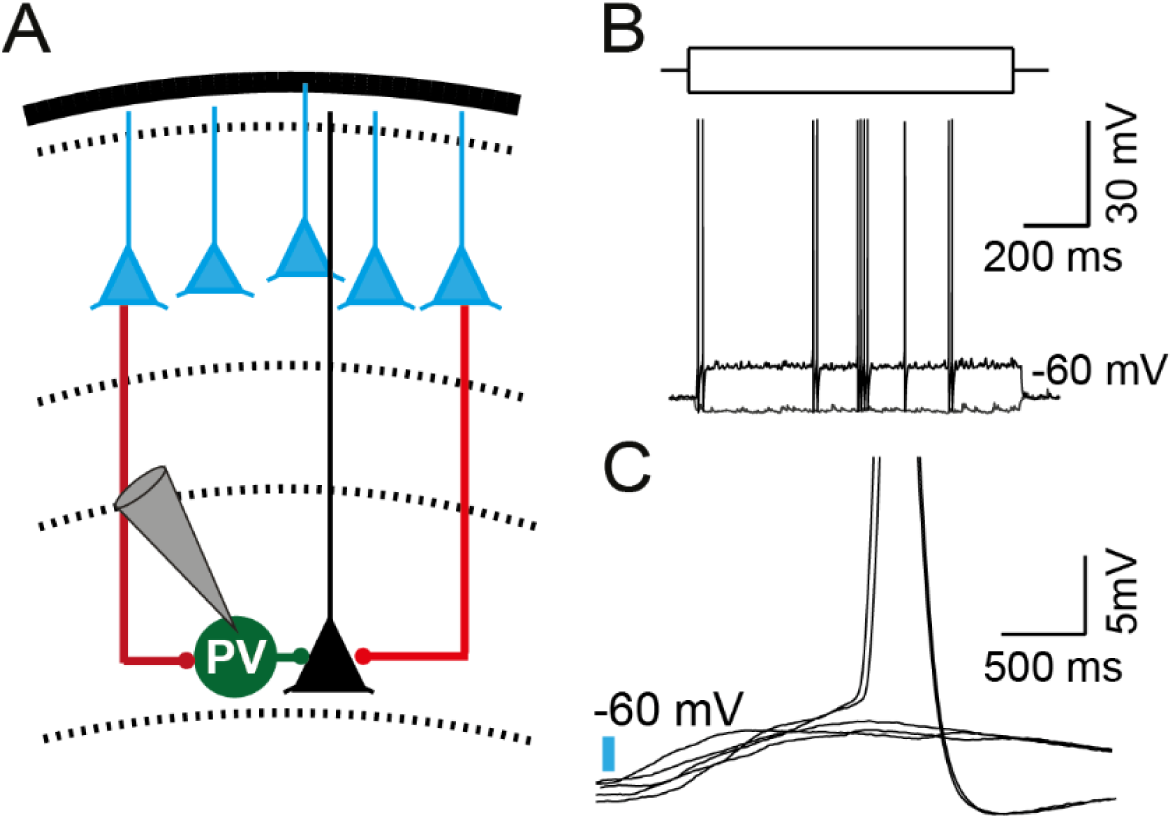
Layer 5 parvalbumin neurons are recruited by photostimulation of layer 2/3 PNs. **(A)** Schematic recording configuration of layer 5 parvalbumin neuron (green circle-PV) stimulated by ChR2+ layer 2/3 PNs (blue triangles). (**B**) Representative traces of a parvalbumin interneuron recorded in current-clamp mode following a hyperpolarizing and depolarizing step of current injection. (**C**) Representative traces of a parvalbumin interneuron recorded in current-clamp mode upon 1ms photostimulation of layer 2/3 ChR2+ PNs. Note the induction of EPSPs above threshold inducing action potentials (action potentials were clipped). 33

**Figure 1-figure supplement 2.**
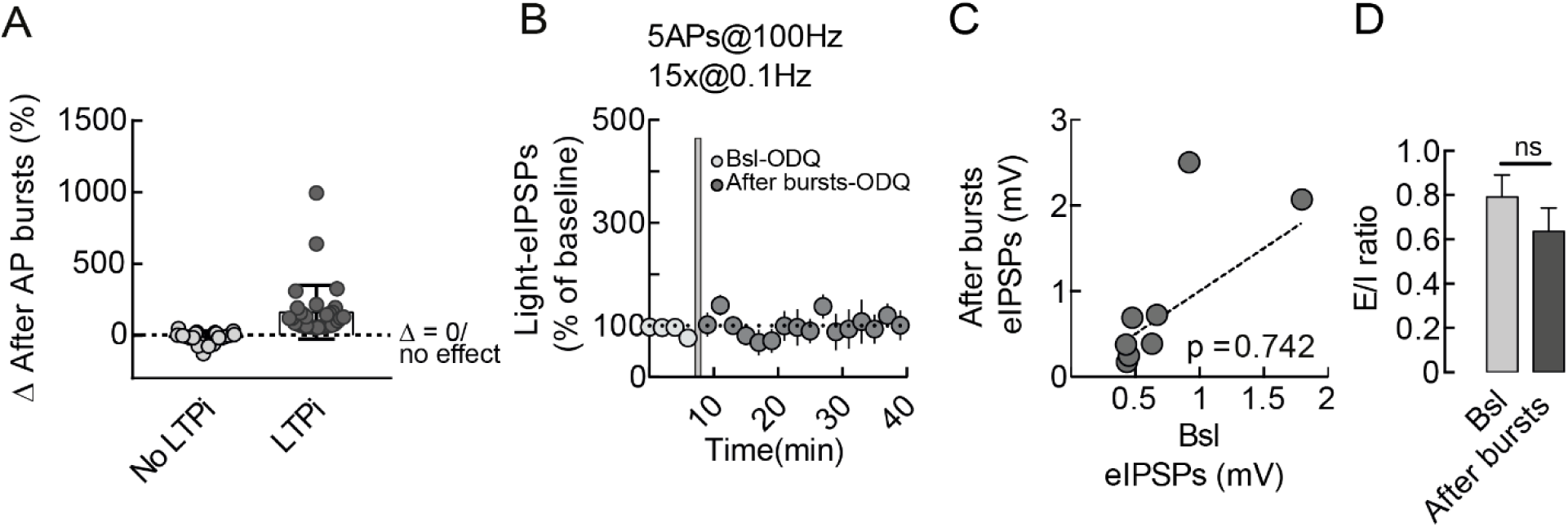
LTPi of feed-forward inhibitory inputs of layer 5 pyramidal neurons is sensitive to nitric oxide signaling. (**A**) Normalized changes of after postsynaptic bursts of action potentials (see Lourenco et al., 2014) from cells in main Figure 1, Figure 2 and Figure 4. Grey light symbols and dark grey symbols refer to pyramidal neurons that did not and that did express LTPi-FFI, respectively. (**B**) Population time courses of normalized light-IPSPs in the presence of the inhibitor of the canonical NO receptor guanylylcyclase (GC) (OQD, 10μM) during baseline (light circles) and after LTPi-FFI induction (black circles) averaged in 2 min. (**C**) Plot of individual eIPSC amplitudes before (x-axes) versus 20 min after postsynaptic bursts (y-axes) in the presence of ODQ. The majority of layer 5 pyramidal neurons failed to express LTPi-FFI. Dotted line indicates unitary values (no change). Error bars indicate SEM. In some cases, the error bars are too small to be visible. (**D**)EPSC/IPSC ratio was not changed upon postsynaptic bursts in the presence of the ODQ(n = 8, p = 0.0781, Wilcoxon matched-pairs signed rank test).

**Figure 2-figure supplement 1.**
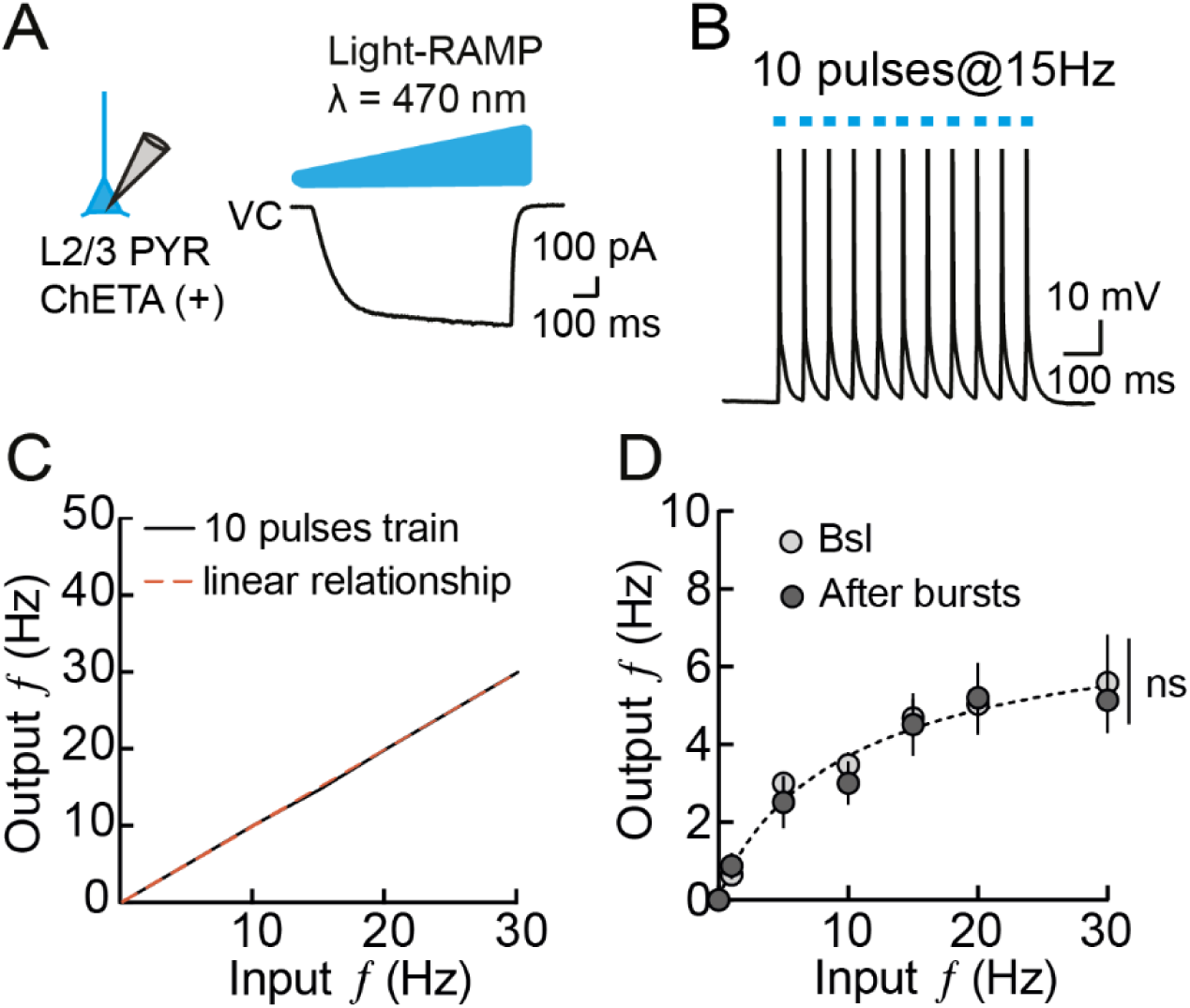
Input/output relationship is constant in cells without LTPi-FFI. (**A**) Schematic recording configuration of a layer 2/3 ChETA+ PNs (blue triangle) and representative trace in voltage-clamp mode upon 1s-long ramp of blue light. (**B**) Representative trace of a layer 2/3 ChETA+ PN following photostimulation of 10 pulses at 15 Hz. Note that each pulse of light induced an action potential (action potentials are clipped). (**C**)Linear output/input relationship of layer 2/3 ChETA+ PNs during input frequencies ranging from 1-30Hz. (**D**) Graph illustrating averaged output firing rate of layer 5 PNs that did not undergo LTPi-FFI. Lines are fits to a Hill function. Note the lack of gain modulation. Error bars indicate SEM.

**Figure 4 supplement 1.**
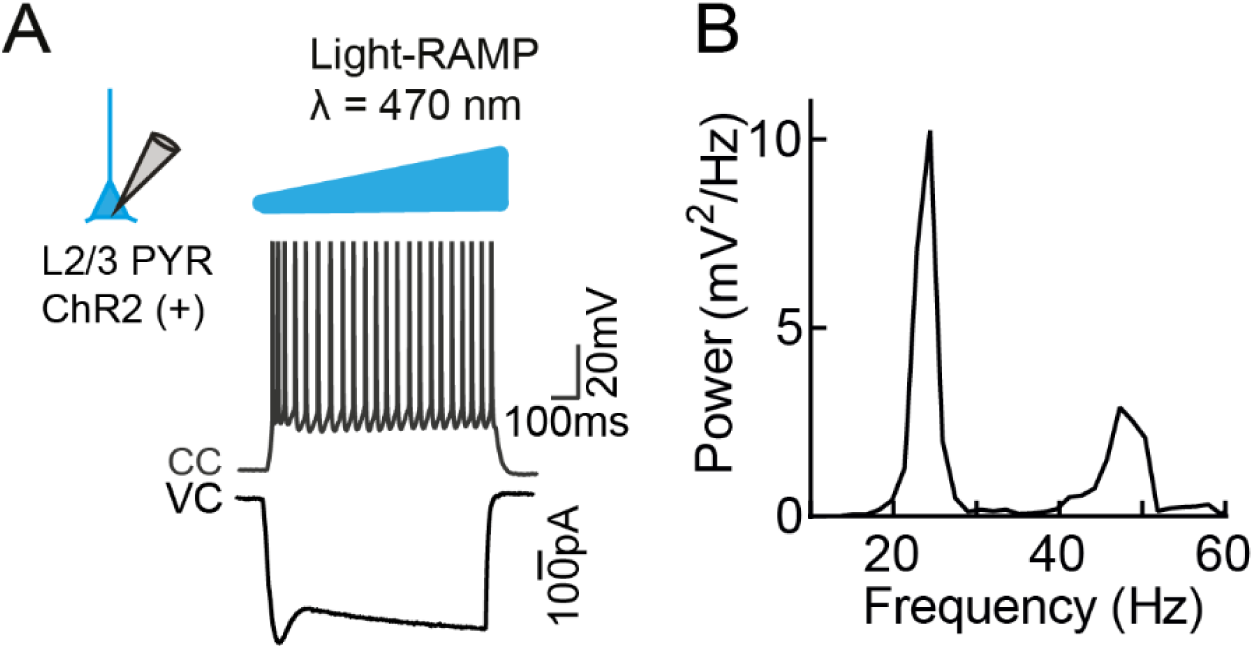
Photostimulation of layer 2/3 ChR2+ PNs induces cortical rhythmic activity. (**A**) Schematic recording configuration of a layer 2/3 ChR2 (+) PNs (blue triangle) and representative trace in voltage-clamp and current-clamp mode upon 1s-long ramp of blue light (action potentials are clipped). (**B**) Power spectrum of action potentials of the cell shown in (**A**). Note the peak of oscillatory activity in the gamma frequency range.

**Figure 5 supplement 1.**
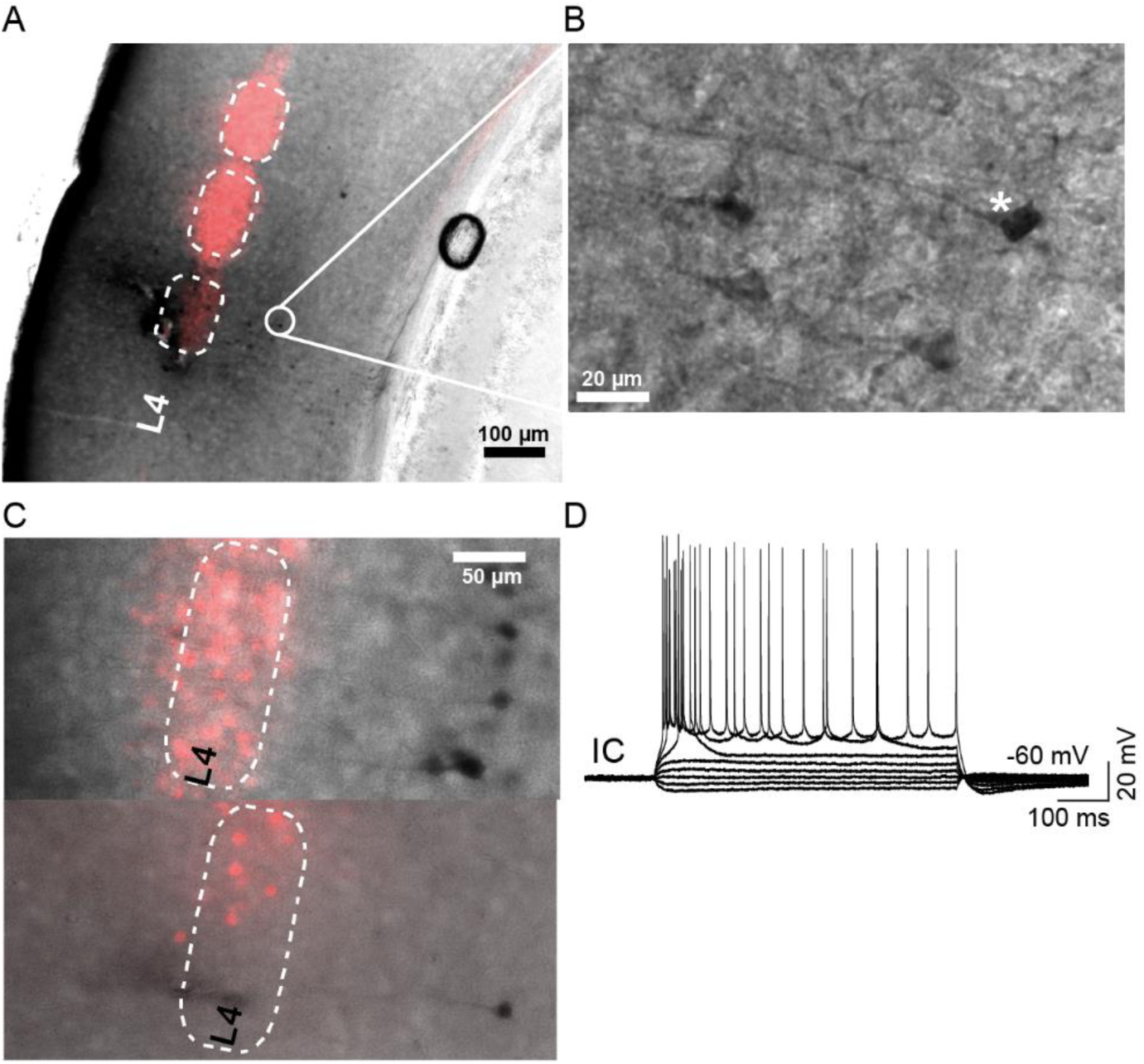
Recording site in layer 5 of S1. (**A**) Photomicrograph of an example coronal section from a Scnn1a x tdTomato mouse following post-hoc processing to detect neurons intracellularly filled by biocytin during *in vivo* recordings. Please note the tdTomato fluorescence highlighting L4. (**B**) inset from (**A**), showing a higher magnification of layer 5 PNs patched in vivo. (**C**) Photomicrographs of two examples showing layer 5 PNs patched in deep layer 5. (**D**) Representative current-clamp (IC) trace from a layer 5 PN in response to a hyperpolarizing and depolarizing current steps. Please note the typical firing behavior of large layer 5 PNs.

**Figure 5 supplement 2.**
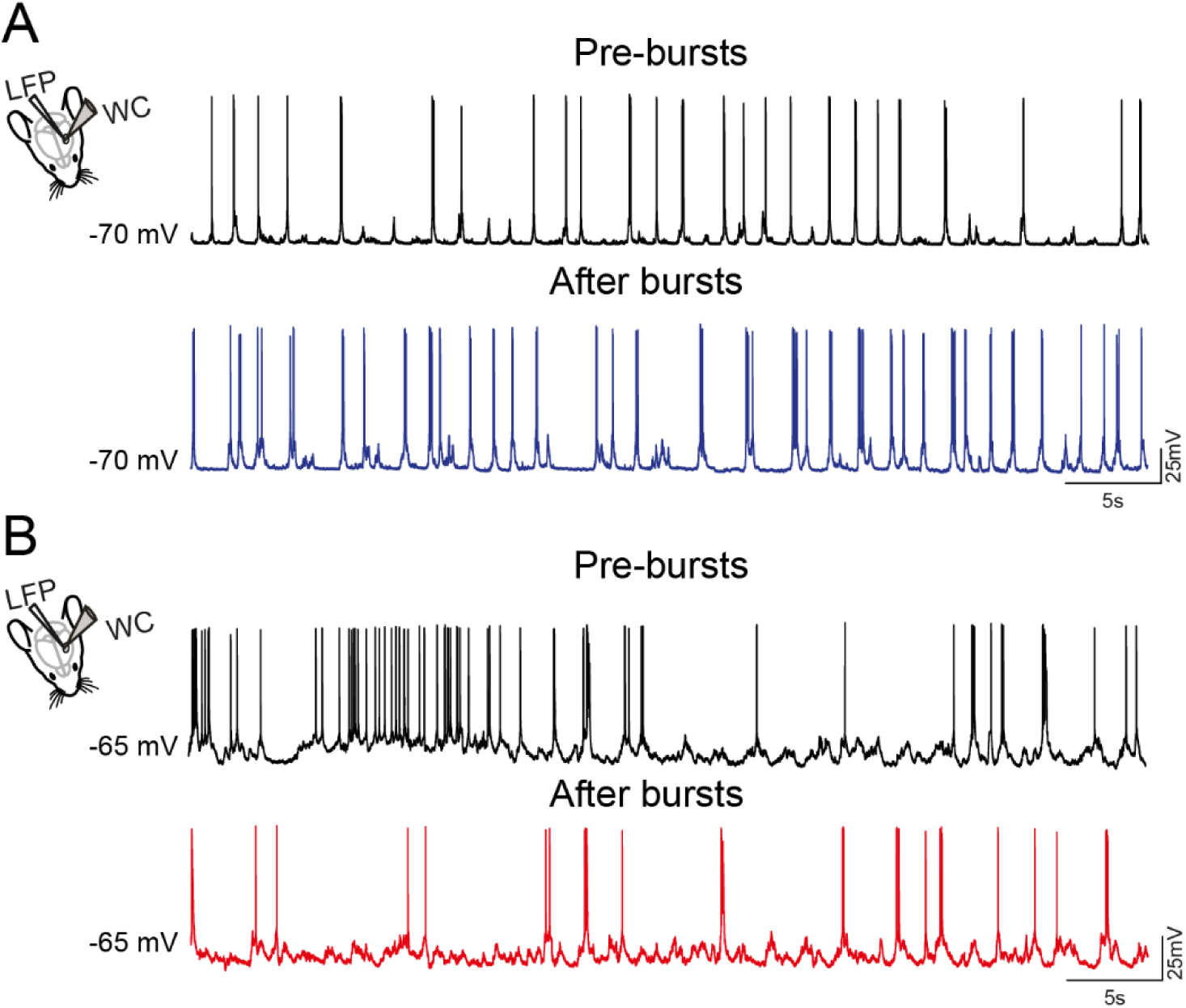
Spontaneous activity of type A and B PNs *in vivo* before and after AP bursts. (**A**) Left panel, Scheme of the recording configuration: LFP and whole-cell (WC) recordings of layer 5 PN in S1, during spontaneous activity. Right: representative current-clamp trace of a type A PN before and after bursts of APs. (**B**) same as in (**A**) but for type B neurons. Please note the increase and decrease in firing in A and B upon postsynaptic bursts activity, respectively.

**Figure 6 supplement 1.**
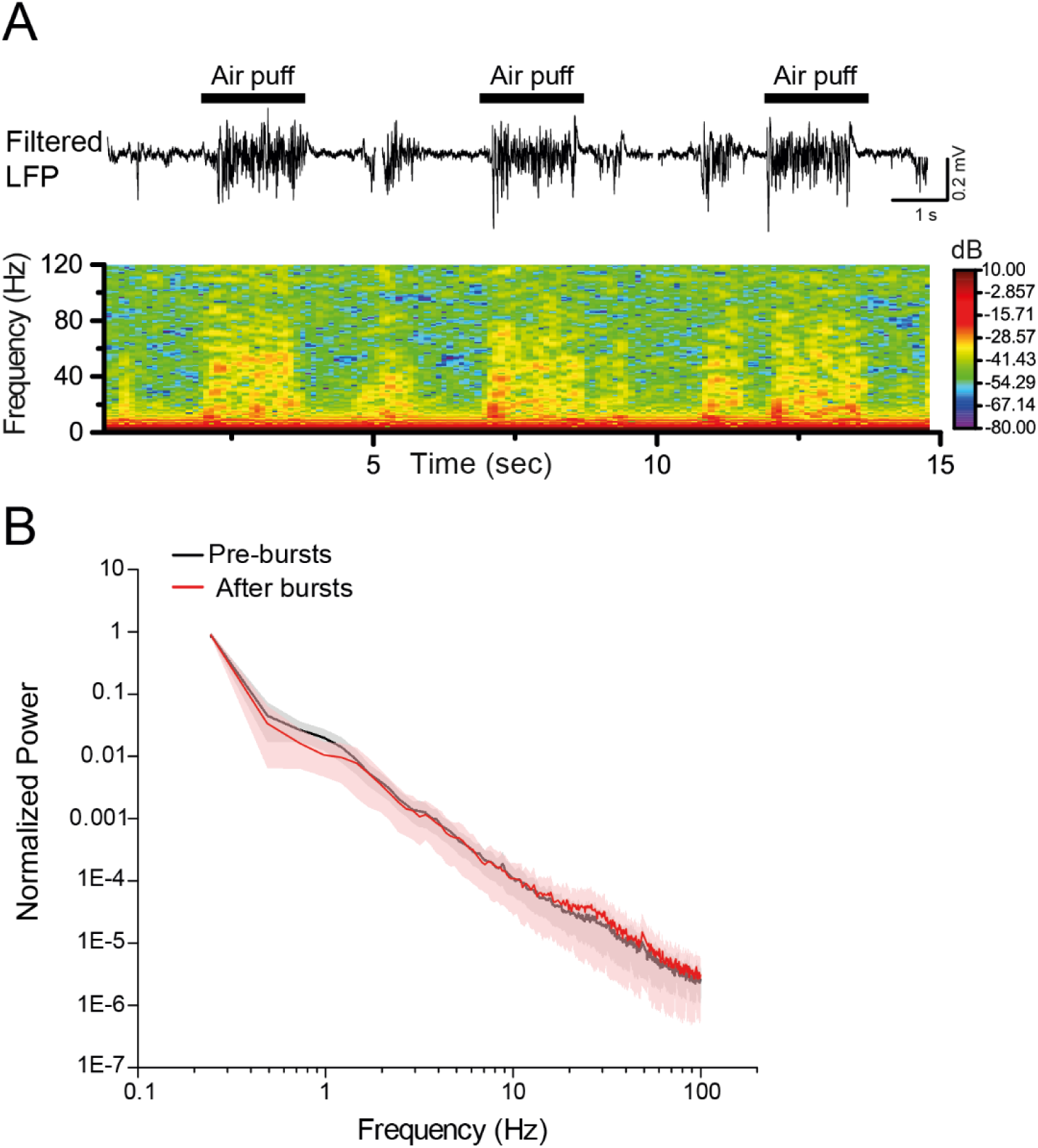
Cortical network dynamics in urethane anesthetized mice. (**A**) Representative trace of filtered LFP (20-100 Hz, top) and the corresponding spectrogram of the unfiltered signal; showing bouts of γ-activity. (**B**) Average normalized LFP power spectra before (black) and after (red) bursts (n = 10 animals).

**Figure 6 supplement 2.**
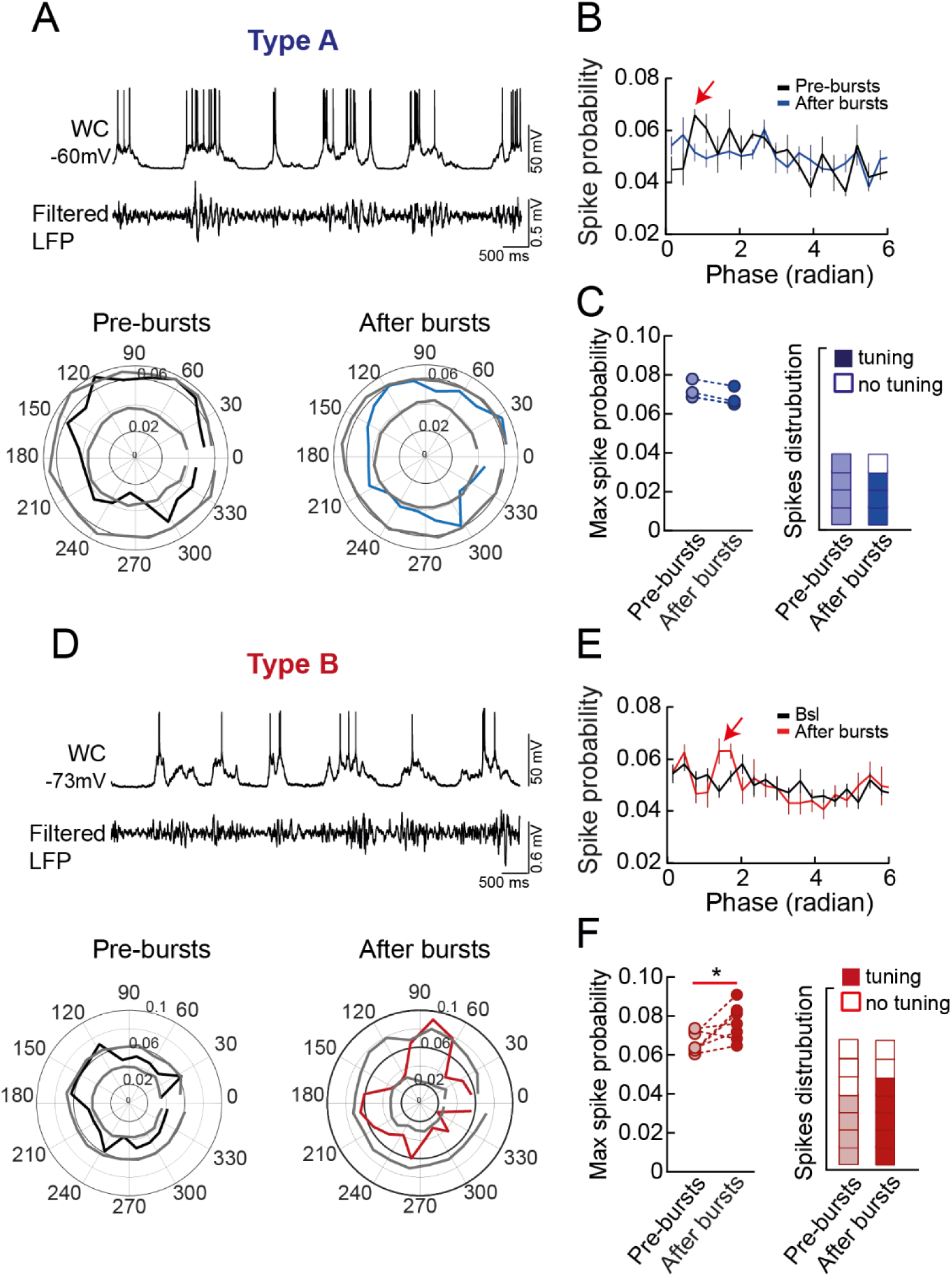
PN spiking activity is poorly tuned to γ-activity non-induced by sensory stimulation, but type B neurons present an increase in max spike probability after bursts. (**A**) Top, representative current-clamp traces showing firing activity in a Type-A layer 5 PN (top trace), and filtered LFP (between 20 and 100 Hz; bottom trace). Bottom: representative polar plots displaying the distribution of APs with γ-phases in the same Type A neuron, before (left, pre-bursts, black trace) and after bursts (right, post-bursts, blue trace). The gray lines correspond to the confidence intervals (set at 1.5 SD threshold) estimated by Monte Carlo methods from 100 independent surrogate data set. (**B**) Population spike distribution across all phases before (black trace) and after bursts (blue trace). Please note the point of maximum (max) spike probability (red arrow). (**C**) Left: summary plot illustrating the max spike probability extracted from each cell before (light blue circles) and after bursts (dark blue circles). Right: plot of the number of cells before (light blue) and after bursts (dark blue) whose spike circular distributions crossed the confidence intervals as shown in A, bottom. White boxes represent non-tuned PNs. (**D**-**F**) same as in (**A**-**C**) but for type B cells. Please note that in these PNs postsynaptic bursts induced a marked increase in max spike probability. In (**B**) and (**E**), the error bars indicate SEM. *p<0.05, Wilcoxon matched-pairs signed rank test.

**Figure 6 supplement 3.**
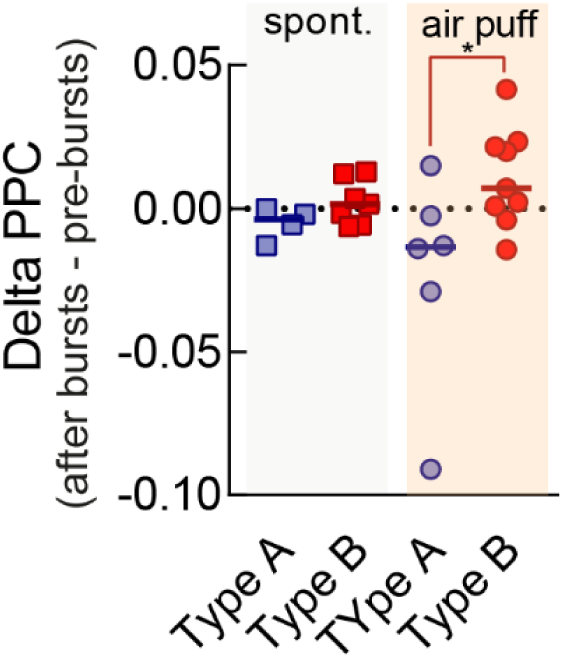
Postsynaptic AP bursts leads to an increase in ΔPPC in type B neurons. Postsynaptic AP bursts leads to a significant increase in phase locking of type B neurons to local rhythmic oscillations in recordings with sensory stimulation as illustrated by ΔPPC (PPC after bursts – PPC baseline) values that are significantly greater than type A neurons.

